# Cell-free biogenesis of bacterial division proto-rings that can constrict liposomes

**DOI:** 10.1101/2020.03.29.009639

**Authors:** Elisa Godino, Jonás Noguera López, Ilias Zarguit, Anne Doerr, Mercedes Jimenez, Germán Rivas, Christophe Danelon

## Abstract

A major challenge towards the realization of an autonomous synthetic cell resides in the encoding of a division machinery in a genetic programme. A key event in the bacterial cell cycle is the assembly of cytoskeletal proteins into a ring that defines the division site. At the onset of the formation of the *Escherichia coli* divisome, a proto-ring consisting of FtsZ and its membrane recruiting proteins takes place. Here, we show that FtsA-FtsZ ring-like structures driven by cell-free gene expression can be reconstituted on planar membranes and inside liposome compartments. Such cytoskeletal structures are found to constrict the membrane and generate budding vesicles, a phenotype that has not been reported before. Additional expression of the FtsZ cross-linker protein ZapA yields more rigid FtsZ bundles that attach to the membrane but fail to produce budding spots or necks in liposomes. These results provide new insights on the self-organization of basic cytoskeletal elements involved in bacterial division. Moreover, they demonstrate that gene-directed protein synthesis and assembly of membrane-constricting FtsZ-rings can be combined in a liposome-based artificial cell.

## INTRODUCTION

Cell-free biology aims at understanding cellular processes by reconstituting biological functions from their isolated elementary components in in vitro model systems. Owing to their openness and easy manipulation, cytoplasmic extracts and systems reconstituted from purified elements are more amenable to customized experimental design and quantitative description compared to living cells. Therefore, the minimal requirements to achieve a particular function can be assessed more reliably. Many complex biological structures and processes taking place in bacterial or eukaryotic cells have already been reconstituted in vitro. Notable achievements include the reconstitution of the minimal translation machinery from *Escherichia coli*^1^, the yeast DNA replication apparatus^2^, filopodial structures^3^, cytoskeleton self-organization and centrosome positioning^4^, egg cytokinesis signaling^5^, DNA segregation with Par^6^, and clathrin-coated buds^7^. Encouraged by the many cellular pieces that have already been reconstituted in vitro, synthetic biologists have now engaged in the construction of an entire cell^8–12^.

One of the hallmarks of living systems is their ability to divide. An obvious starting point to conceiving a biology-inspired division mechanism in artificial cells is to consider the canonical pathways taking place in prokaryotes. In most bacteria, symmetrical cell division proceeds by forming a constriction ring that eventually splits the mother cell into equally sized daughter cells^13^. At an earlier stage of cytokinesis, a proto-ring composed of the FtsZ, FtsA and ZipA proteins, assembles on the inner leaflet of the cytoplasmic membrane at the future division site^14–16^. The tubulin-related FtsZ is the core constituent of the proto-ring. FtsZ is a GTPase that can polymerize into protofilaments^17,18^. Anchoring of FtsZ protofilaments to the cytoplasmic membrane is mediated by ZipA and the actin homologue FtsA^19–21^. This process is regulated by accessory proteins belonging to the Zap family^22^.

Earlier attempts to divide cell-like liposomal compartments have focused on the reconstitution of the Z-ring from purified proteins^23,24^. These studies have shown that FtsZ aided by one of its anchoring protein partners - or a chimeric FtsZ bearing a membrane targeting segment^25^ - can self-organize into filament patterns on supported lipid membranes^25–28^. When encapsulated inside vesicles, the elementary cytoskeletal proteins form ring-like structures that can deform the liposome membrane^24,29,30^. Whether FtsZ filaments alone exert a contractile force and contribute to the final stage of division remains a subject of debate^31,32^ and evidence for complete liposome division is still lacking.

A conceptual issue which is inherent to reconstitution assays solely relying on purified proteins, is the impossibility to maintain steady amounts of cytoskeletal proteins from internal mechanisms as the compartment undergoes division. Another problem raised by conventional cell-free assays is the use of oversimplified buffer compositions that have been tailored for a particular set of enzymatic reactions but fail to reproduce the cytoplasmic environment.

Herein, we address these issues by encoding *E. coli* division proteins on DNA templates. Genetic control over protein production offers a general solution to achieve self-replication, as well as self-regulation by establishing feedback loops. In this context, the PURE system, a minimal gene expression system reconstituted primarily from *E. coli* constituents^1^ was employed. Different types of proteins and biological functions have already been synthesized de novo with the PURE system, including membrane-associated proteins^10–12,33^. Moreover, by containing all relevant factors for gene expression, the PURE system emulates more closely the molecular composition of the bacterial cytoplasm than simple buffers. In the present study we utilized PURE*frex*2.0 which provides the best combination of protein yield and expression lifespan^34,35^.

We show that cell-free expressed FtsA is able to recruit FtsZ polymers, forming large-scale two-dimensional networks of curved and ring-like structures in the absence of bundling factors. When the entire set of reactions is encapsulated inside liposomes, proto-rings of FtsA-FtsZ filaments are found to constrict the membrane and generate budding vesicles. Co-expression of ZapA, a native stabilizer of FtsZ filaments, yields stiffer FtsZ bundles attached to the membrane that fail to constrict into bud necks. FtsZ cytosekeltal structures are also investigated with ZipA membrane-anchor protein. We find that in our low-volume supported lipid bilayers (SLBs) assays with ZipA and ≤ 3 µM FtsZ, the generic crowding agent Ficoll70 is necessary to elicit bundle formation. Cell-free expressed ZapA obviates the need of Ficoll70 and promotes formation of cytoskeletal networks with different, likely more physiological, morphology and protein monomer dynamics. The prospects of further improvement suggest that the DNA-programmed hierarchical assembly of the Z-ring in liposomes is a promising strategy for dividing synthetic cells. In addition, our approach to reconstituting cellular processes in PURE system provides a generic platform that fills the gap between classical in vitro and in cellulo experiments.

## RESULTS

### Cell-free synthesized FtsA drives the formation of curved FtsZ filaments

An essential component of the *E. coli* division proto-ring is FtsA, a homologue of actin that anchors FtsZ filaments to the cytoplasmic membrane. To bypass the difficult purification of FtsA^28^, we directly expressed a sequence optimized *ftsA*_opt_ gene on a supported lipid bilayer (SLB) (**Fig. 1a**). In the presence of 3 µM purified FtsZ-A647, curved filaments and ring-like structures formed on the membrane (**Fig. 1b, Supplementary Fig. 1**), concurring with previous reports^12,28^.

**Figure 1:**
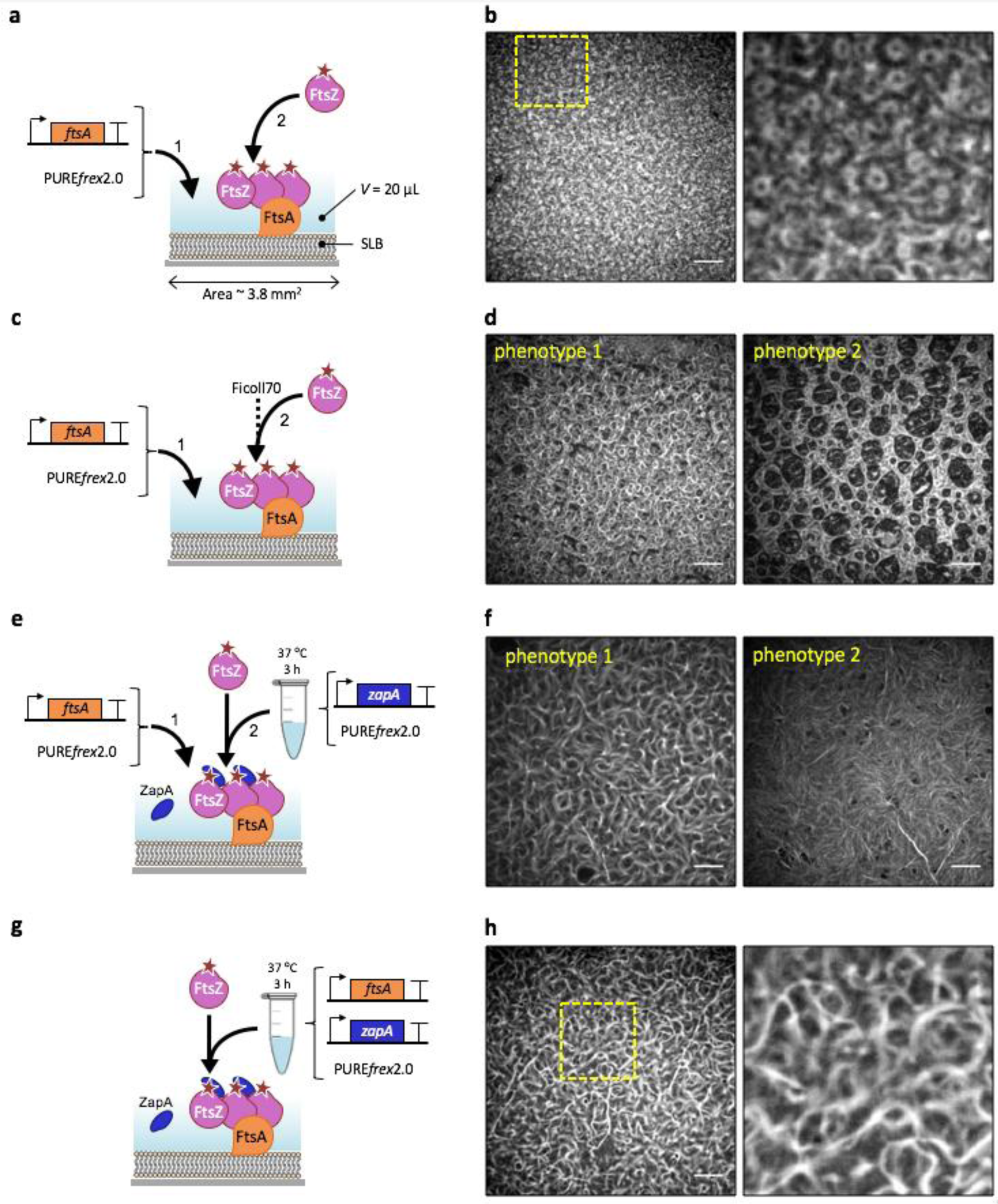
Cell-free expressed FtsA recruits FtsZ to an SLB and drives the formation of ring-like structures. **a**, Schematic representation of the SLB assays with FtsA directly expressed on the membrane. Purified FtsZ-A647 (3 µM) was added. The sequence-optimized construct *ftsA*_opt_ was used. **b**, Fluorescence image of FtsZ-A647 forming curved filaments and rings in the presence of in situ synthesized FtsA. The zoom-in image (right) corresponds to the framed region in the left image. **c**, As in **a**, but the solution was supplemented with Ficoll70. **d**, Fluorescence image of FtsZ-A647 forming curved filaments and rings in the presence of in situ synthesized FtsA and Ficoll70. Two representative filament network morphologies are shown. **e**, Schematic illustration of the SLB assays with separately expressed FtsA and ZapA. The constructs *ftsA*_opt_ and *zapA* were used. Purified FtsZ-A647 (3 µM) was included. **f**, Fluorescence images of FtsZ-A647 displaying two representative phenotypes from different regions of an SLB. **g**, Schematic illustration of the SLB assays with co-expressed FtsA and ZapA from *ftsA*_opt_ and *zapA*_opt_ constructs. Purified FtsZ-A647 (3 µM) was added. **h**, Fluorescence images of FtsZ-A647. The zoom-in image (right) corresponds to the framed area in the left image. Scale bars indicate 10 µm.

Promoting lateral interactions of FtsZ protofilaments stimulates the formation of higher-order cytoskeletal structures in vitro^36^. However, little is known about how the nature of these lateral interactions influences the morphology of the FtsZ network. Therefore, we decided to investigate the architecture and dynamics of FtsZ protofilaments in a molecular environment that favours lateral interactions. First, we employed Ficoll70, a generic crowding agent known to elicit FtsZ bundle formation (**Fig. 1c**). Large SLB areas were covered with curved filaments, rings of different sizes (most having a diameter of 1 – 2 µm) and large circular patterns (**Fig. 1d**).

Although Ficoll70 is commonly used as a macromolecular crowder to mimic cytoplasmic conditions^24,33,37,38^, we reasoned that ZapA, an in vivo regulator of FtsZ polymerization, would provide a more targeted and native mechanism to elicit lateral interaction, thus conferring physiologically relevant properties of cytoskeletal patterns. For this reason, ZapA was produced in PURE*frex*2.0 starting from its native gene sequence. Substitution of Ficoll70 with cell-free synthesized ZapA produced curved bundles but also long and straight filaments of FtsZ-A647 recruited to the membrane by cell-free synthesized FtsA (**Fig. 1e,f**). Challenging PURE system to co-express both *ftsA*_opt_ and *zapA*_opt_ genes in a single reaction led also to the formation of bended and straight bundles (**Fig. 1g,h, Supplementary Fig. 2**). Both the curvature and occurrence of branching points of the filament network were reduced in the presence of Ficoll70 or ZapA (**Supplementary Fig. 3**). Despite morphological differences observed in the protein patterns between the Ficoll70- and ZapA-containing samples, quantitative analysis of FtsZ subunit dynamics by fluorescence recovery after photobleaching (FRAP) revealed similar recovery halftime values under the tested conditions (**Supplementary Fig. 4)**.

### Bundling of FtsZ filaments is required for the formation of long sZipA-FtsZ cytoskeletal structures in low-volume SLB assays

We then examined the self-organization of FtsZ with the membrane anchor soluble ZipA (sZipA) using purified proteins supplied in a minimal buffer or in PURE*frex*2.0 background. We found that in our low-volume SLB assays with ≤ 3 µM FtsZ, filament bundling with either Ficoll70 or ZapA was required to trigger large-scale cytoskeletal networks (**Fig. 2a,b, Supplementary Note 1**). This result contrasts with previous observations^20,28^, highlighting the role of the total reaction volume as a control parameter in the formation of cytoskeletal patterns. The network morphology, as well as the sZipA and FtsZ monomer dynamics differ whether lateral interactions are promoted by the artificial molecular crowder Ficoll70 or by cell-free synthesized ZapA (**Fig. 2c-f, Supplementary Fig. 5-8, 10**). Moreover, we found that increasing the expression level of ZapA results in more stable filaments that can even extend above the SLB (**Supplementary Fig. 6,7,9**). Taken together, our results show that ZapA encourages the formation of membrane-bound sZipA-FtsZ filament network having a different - presumably more physiological - morphology and subunit turnover compared to Ficoll70.

**Figure 2:**
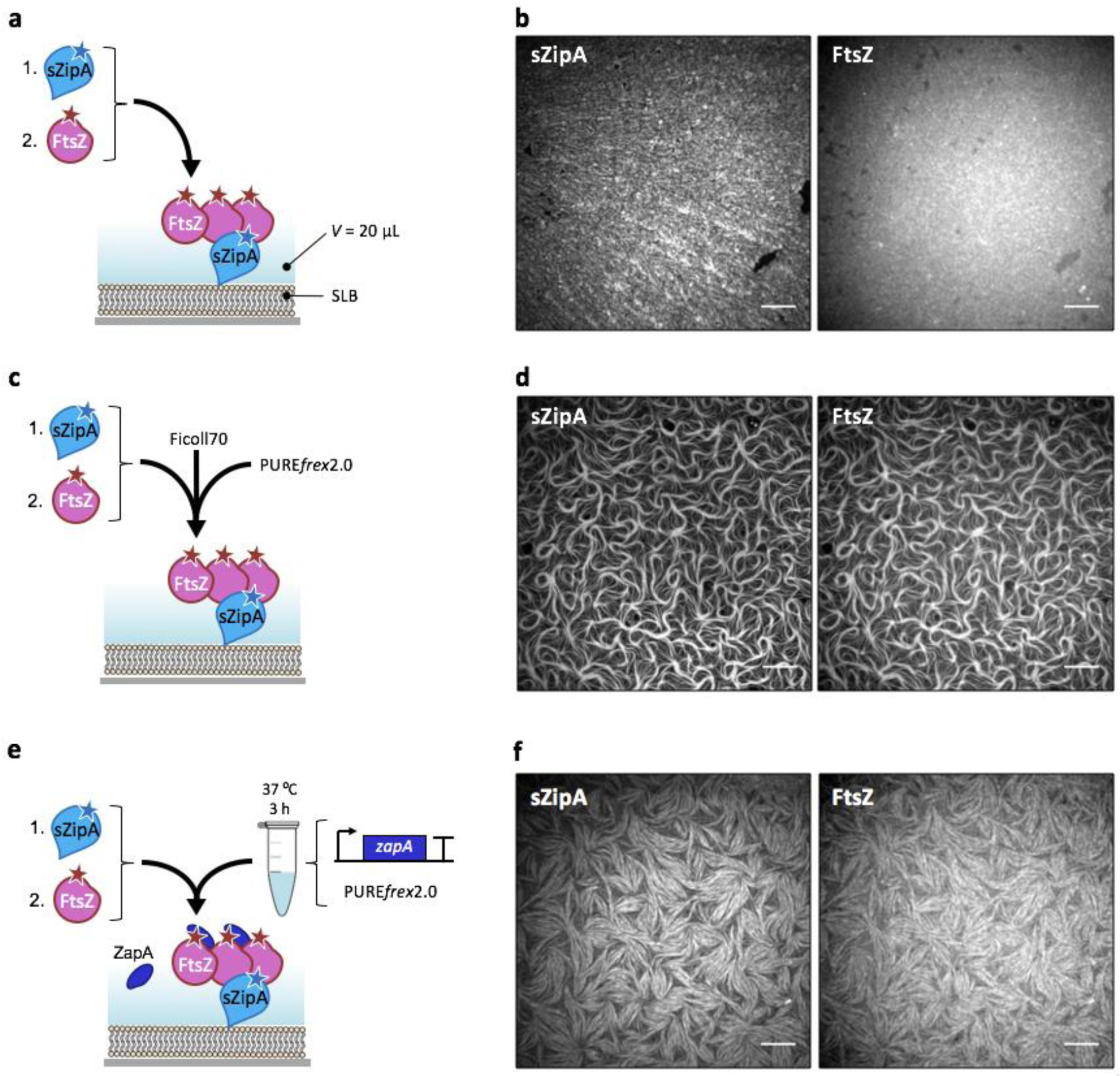
Purified sZipA and FtsZ form co-filament networks in the PURE system. **a**, Schematic representation of the SLB assays. Purified sZipA-A488 was first incubated on an SLB. The solution on top of the SLB was replaced by a minimal reaction buffer containing 3 µM purified FtsZ-A647 and 2 mM GTP. **b**, Fluorescence images of sZipA-A488 (left) and FtsZ-A647 (right) in the minimal reaction buffer without Ficoll70. **c**, Schematic representation of the SLB assays. Purified sZipA-A488 was first incubated on an SLB. The solution on top of the SLB was replaced by PURE*frex*2.0 supplemented with 3 µM purified FtsZ-A647, 2 mM GTP and 50 g L^−1^ Ficoll70. **d**, Fluorescence images of sZipA-A488 (left) and FtsZ-A647 (right) in PURE*frex*2.0 with Ficoll70. Large-scale filaments with co-localizing FtsZ-A647 and sZipA-A488 are exclusively observed in the presence of Ficoll70. This conclusion is valid in both the minimal reaction buffer and in PURE*frex*2.0 background. More fields of view are displayed in **Supplementary Fig. 5. e**, Schematic illustration of the SLB assays with purified FtsZ-A647 (3 µM) and cell-free synthesized ZapA incubated on top of an sZipA-A488-bound SLB. ZapA was expressed from the native gene *zapA*. **f**, Fluorescence images of sZipA-A488 (left) and FtsZ-A647 (right) in a sample containing cell-free synthesized ZapA and additional 2 mM GTP. Different cytoskeletal network phenotypes were observed when ZapA concentration was increased upon expression of the optimized *zapA*_opt_ construct (**Supplementary Fig. 7**). Scale bars indicate 10 µm.

We conclude from these SLB experiments that short, curved filaments and rings that resemble physiological structures are more prominent with FtsA and can develop in the absence of a bundling agent compared to sZipA.

### FtsZ and internally synthesized FtsA self-organize into ring-like structures that constrict liposomes

The identification of FtsZ and FtsA as the minimal molecular set to obtain membrane-anchored curved filaments and rings in the PURE system prompted us to reconstitute FtsA-FtsZ cytoskeletal networks inside liposomes (**Fig. 3a**). The cell-free gene expression solution was supplemented with adenosine triphosphate (ATP, additional 2 mM), guanosine triphosphate (GTP, additional 2 mM) and a mixture of highly purified chaperones (DnaK mix). Although energy regeneration components are present in the PURE system, extra ATP and GTP were provided to compensate for the extra demand from FtsA and FtsZ. Purified FtsZ-A647 was used to visualize protein localization by laser scanning confocal microscopy. FtsZ-A647 was employed at 3 µM concentration which is similar to that measured *in vivo* (∼3.5 µM)^39^. Liposomes were formed by natural swelling, with a composition of zwitterionic PC and PE phospholipids, anionic PG and cardiolipin, and a small fraction of TexasRed-conjugated lipid for membrane imaging^35^. Such a lipid mixture and liposome preparation method have proved compatible with the cell-free synthesis of membrane-associated enzymes^10^, DNA replication proteins^11^ and division proteins^12^. Liposome size distribution ranges from ∼1 µm up to over 15 µm in diameter, which provides a more relevant bacterial cell-size compartment than >20 µm liposomes. In contrast with previous studies^33^, no crowding agent was included during liposome formation. Actually, we found that Ficoll70 impairs formation of gene-expressing liposomes with our methodology (**Supplementary Fig. 11**).

**Figure 3:**
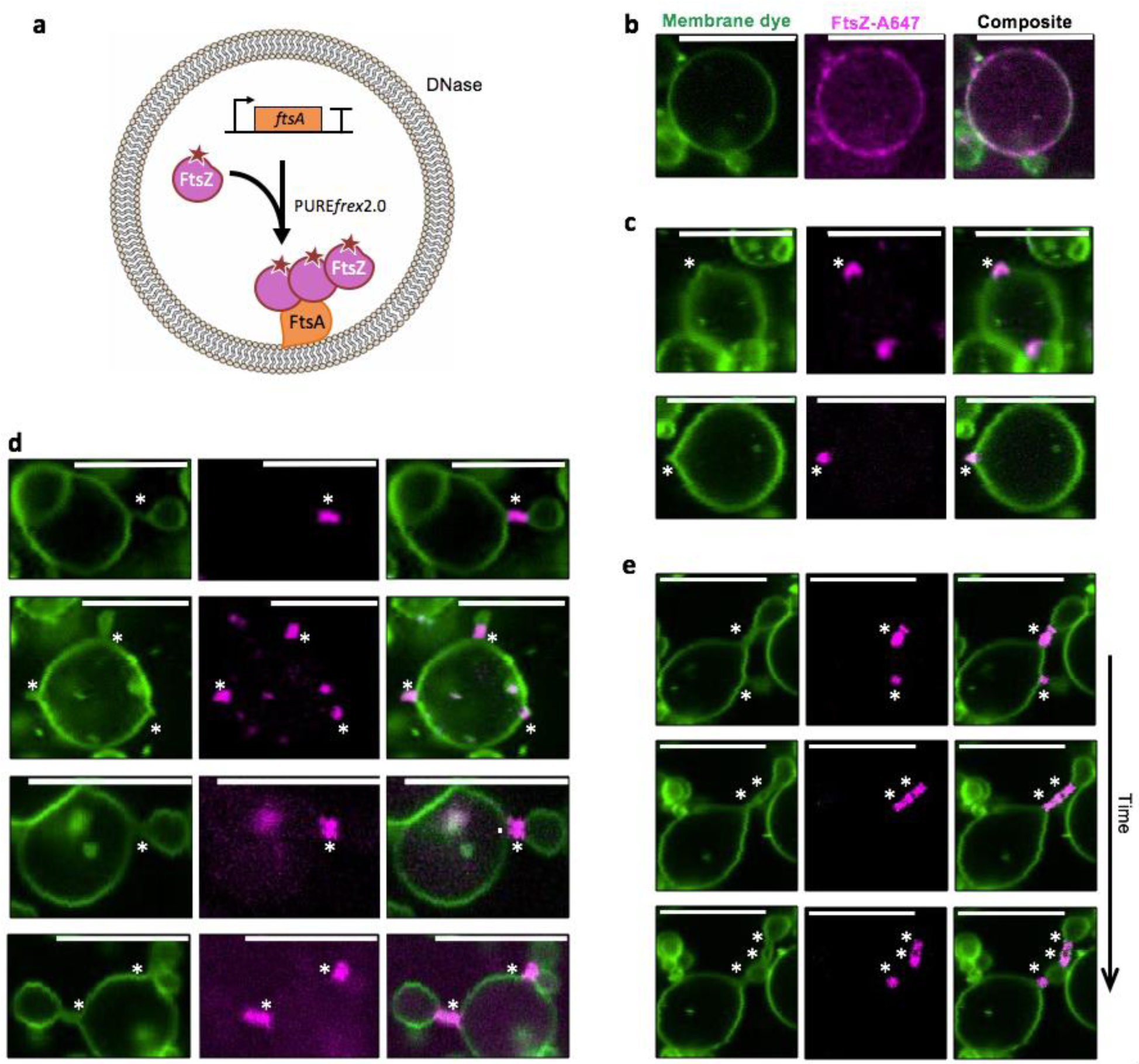
In-liposome synthesized FtsA assembles with FtsZ into ring-like structures that drive vesicle budding. **a**, Schematic illustration of liposome reconstitution assays. The *ftsA*_opt_ DNA template was expressed within phospholipid vesicles in the presence of 3 µM purified FtsZ-A647. **b**-**d**, Confocal fluorescence images of liposomes exhibiting different morphologies of FtsZ-FtsA cytoskeletal structures and membrane remodelling: recruitment of proteins to the membrane in the form of clusters with no visible membrane deformation (**b**), budding spots induced by local accumulation of FtsZ-FtsA (**c**), and budding vesicles from a parental liposome with a clear FtsA-FtsZ-coated membrane neck (**d**). **e**, Time series images showing that a ring-forming protein cluster localized at a constriction site can split, which induces multiple necks separated by blebbing vesicles (see **Movie 1**). Fluorescence from the membrane dye is coloured in green and FtsZ-A647 signal is in magenta. The composite image is the overlay of the two channels. Asterisks indicate budding spots or constriction sites. Scale bars represent 10 µm. More examples of liposomes are shown in **Supplementary Fig. 13** and **14**.

In control experiments where the *ftsA*_opt_ gene was omitted, FtsZ was exclusively located in the liposome lumen **(Supplementary Fig. 12**). De novo synthesized FtsA successfully recruited FtsZ on the membrane as shown by the co-localization of the FtsZ-A647 and membrane dye signals (**Fig. 3, Supplementary Fig. 13**). Although homogeneous recruitment of FtsZ to the membrane was commonly found within the liposome population, the majority of the liposomes displayed regions with patches of FtsZ on the inner surface of the membrane (**Fig. 3b**). Noticeably, the membrane spots with clustered FtsZ coincide with different types of membrane remodelling. In some cases, the recruited FtsZ localizes with outward membrane deformation or short protrusions (**Fig. 3c, Supplementary Fig. 14**). In other instances, the protrusions developed into vesicles or blebs tethered to the parental liposome through a membrane neck coated with FtsZ (**Fig. 3d, Supplementary Fig. 14**). Sometimes, the budding neck extends over a few microns in the form of a tubular structure containing one or a few FtsA-FtsZ rings (**Fig. 3d,e**). Interestingly, these blebbing structures are dynamic. Events, such as appearance of new constriction sites, growing vesicles and diffusion of protein rings along the tube axis were observed (**Fig. 3e, Movie 1**). Although membrane recruitment of FtsZ in the form of patches was visible already within 2 h of expression, major liposome remodelling events, such as budding spots and elongated blebs were observed only after 3 to 4 h. Moreover, after 6 h expression, small vesicles were found to agglutinate to larger liposomes (**Supplementary Fig. 12**). It is unclear whether the FtsZ-coated membrane necks can close to release mature vesicles. Therefore, we cannot ascertain that these small vesicles are reminiscent to division events. Yet, these aggregated vesicles were not observed when the *ftsA*_opt_ gene was omitted (**Supplementary Fig. 12**), indicating that this global remodelling is dependent on the expression of FtsA.

We then decided to investigate how the presence of cell-free expressed ZapA would modulate the properties of the cytoskeletal patterns in liposomes (**Fig. 4a**). Co-expression of *ftsA*_opt_ and *zapA*_opt_ DNA constructs induced formation of FtsZ-A647 clusters on the inner surface of the membrane (**Fig. 4b**). Liposomes with different cytoskeletal protein phenotypes were observed, such as homogeneous coating to the membrane, patches or filaments, and large ring-like structures. Bundles of FtsZ polymers adopting apparent ring-like structures predominantly localize at the interface of two liposomes, coinciding with a membrane septum (**Fig. 4b**). However, ZapA abolishes the formation of membrane protrusions, vesicle budding and clustering of FtsZ on tubular membrane structures. These results indicate that the mechanical properties of FtsZ-ZapA bundles impede the formation of membrane-constricting, high-curvature cytoskeletal filaments, which suggests that temporal regulation of the local concentration of ZapA might play a role in the constriction of the FtZ ring.

**Figure 4:**
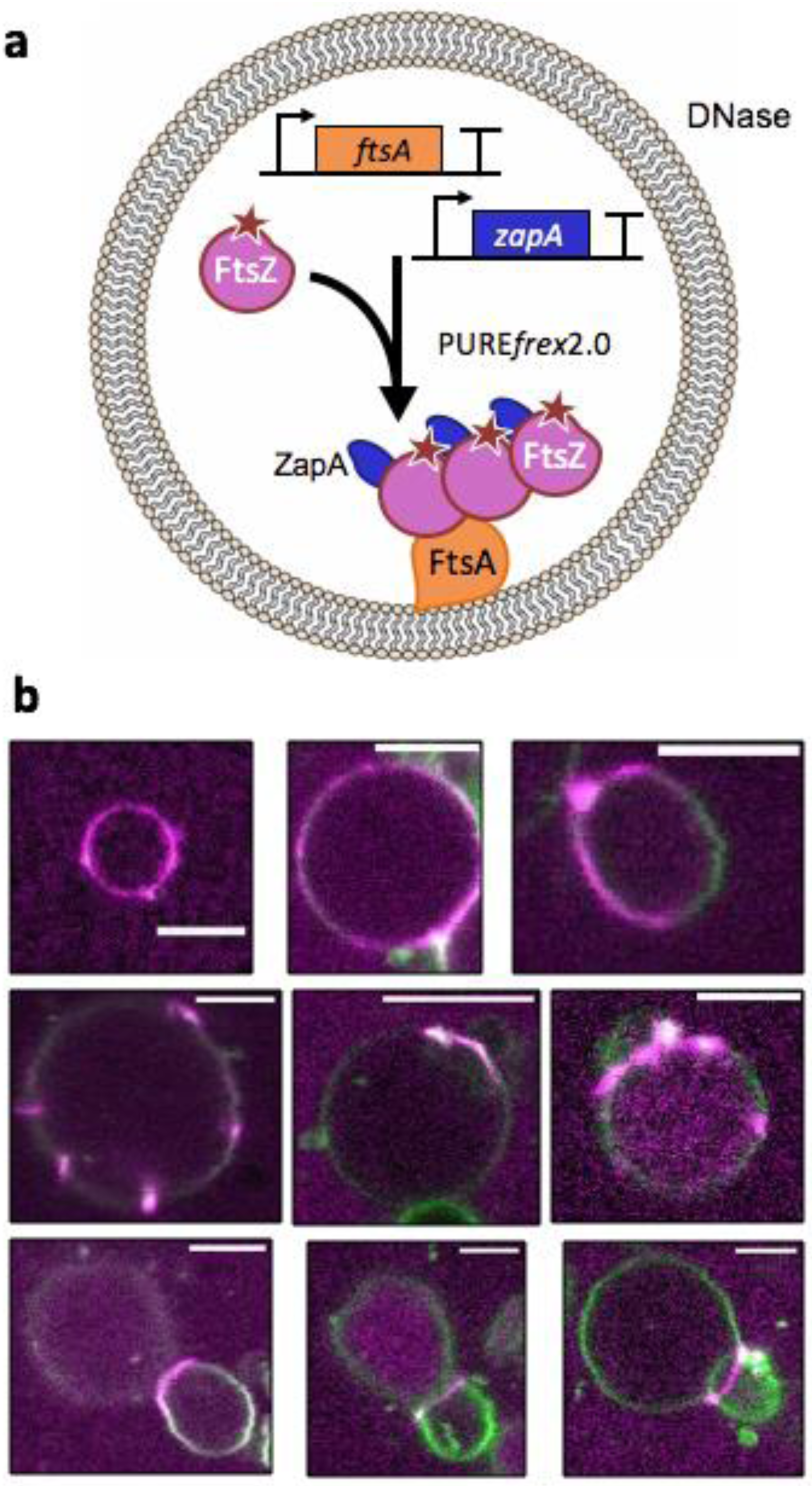
Co-expressed FtsA and ZapA organize FtsZ into long membrane-tethered bundles within liposomes. **a**, Schematic illustration of liposome reconstitution assays with 3 µM purified FtsZ-A647 and co-expression of the *ftsA*_opt_ and *zapA*_opt_ DNA constructs. **b**, Confocal fluorescence images of liposomes exhibiting membrane recruitment of FtsZ-A647 after 3 h incubation. Fluorescence from the membrane dye is coloured in green and FtsZ-A647 signal is in magenta. Only the composite images are displayed. Scale bars represent 5 µm.

Collectively, the results demonstrate that gene-based reconstitution of membrane-constricting cytoskeletal protein filaments within liposomes is feasible. Moreover, FtsA and FtsZ form the minimal architecture to establish E. coli cell division proto-rings from native constituens.

## DISCUSSION

Compartmentalization of PURE system and of the Z-ring constituents inside liposomes provides a realistic cellular environment. Purified FtsZ and FtsA proteins have already been enclosed within small (diameter < 200 nm) liposomes^30^ or giant vesicles^40^. In these earlier studies, membrane-tethered protofilaments of FtsZ could assemble with FtsA*, a mutant of FtsA that cannot polymerize^41,42^. In another report, the FtsZ-sfGFP fusion protein was recruited to the membrane of giant liposomes (diameter 15 – 100 µm) by FtsA in the presence of Ficoll70^33^. In their work, Furusato et al. reported a homogenous recruitment of FtsZ to the membrane in the presence of FtsA but no membrane deformation^33^. Local reshaping of liposomes was exclusively observed with ZipA as the FtsZ membrane anchor, but no constriction sites nor protein ring-like structures were observed^33^. Here, we show that wild-type FtsA and FtsZ are capable to deform the membrane in PURE system-loaded liposomes with a size < 15 µm. The FtsA-dependent FtsZ recruitment on the membrane frequently induces the formation of FtsZ clusters that constrict the liposome membrane into bud necks. It is clear from our data that FtsZ, assisted by FtsA, does not preferentially accommodate to pre-existing membrane areas with a high curvature. Conversely, membrane constriction in the form of a narrow neck connecting the mother and budding vesicles results from FtsA-FtsZ clustering. Noteworthily, this type of membrane remodelling and cytoskeletal protein organization was not observed in previous reports^23,24,33,43^.

Not every liposome exhibits the same phenotype with regard to FtsZ recruitment and membrane deformation. This disparity is presumably the manifestation of the probabilistic encapsulation of all PURE system components and DNA, which leads to a large heterogeneity in FtsA expression levels, as recently quantified with a fluorescence reporter gene^35^. It is therefore difficult to know the precise concentration of synthesized FtsA in individual liposomes and to correlate it with a particular phenotype.

Further investigations will be necessary for directing the assembly of an all-gene based contractile FtsZ proto-ring that can divide liposomes. Although we do not exclude that assisting proteins, such as the Min system^12^ and the FtsZ-interacting partners SlmA^44^ and ZapB^45^, might have to be introduced, the present results suggest that expression of FtsA and FtsZ might suffice to generate daughter vesicles of a few microns in size. The precise timing of protein interaction is essential for the hierarchical assembly of the proto-ring. This represents a major issue that is inherent to in-liposome compartmentalization of purified cytoskeletal proteins or with temporally unregulated expression of multiple genes. An additional level of temporal control that might be decisive for sequential assembly of the Z-ring constituents could be provided by regulating the expression of individual genes through transcriptional circuits, such as cascade or feedback motifs^46^. Mindful of the limitations to apprehend the PURE system^34^ and to rationally design liposomes harbouring desirable properties encoded in genes, we believe that in vitro evolutionary optimization, by exploring a wide genetic diversity, provides additional opportunities to build cellular functions, and FtsZ proto-rings in particular.

## METHODS

### DNA constructs

*ftsZ* and *ftsA* gene fragments were amplified by standard polymerase chain reaction (PCR) from the chromosomal *E. coli* BL21 DNA with primers 509 and 374 (*ftsZ*), and 508 and 376 (*ftsA*) (**Supplementary Table 1**). These primers contain overhangs for Gibson assembly with the pET11-a plasmid. PCR products were checked on a 1% agarose gel stained with EtBr or SYBR safe, imaged with a ChemiDocTM Imaging System (BioRad Laboratories), and purified with the Wizard SV Gel kit (Promega). The purified DNA was incubated with DpnI (New England BioLabs®, Inc.) to remove residual plasmid and the linear DNA was purified again with Wizard SV Gel kit. DNA concentration and purity were measured using a ND-1000 UV-Vis Spectrophotometer (Nanodrop Technologies). Gibson assembly (Gibson Assembly® Master Mix of New England BioLabs®, Inc.) was performed at equimolar concentrations of linearized plasmid (pET11-a) and DNA fragments for 1 h at 50 °C. *E. coli* TOP10 competent cells were transformed with the Gibson assembly products by heat shock. Cells were centrifuged, resuspended in 50 µL of fresh prechilled liquid lysogen broth (LB) medium and incubated for 1 h at 37 °C and 250 rpm. The cultures were plated on solid LB medium with ampicillin and grew overnight at 37 °C. Colonies were picked up and cultured in 1 mL of liquid LB medium with 50 µg µL^−1^ of ampicillin in 1.5-mL Eppendorf tubes for 6 h at 37 °C and 250 rpm. Plasmid purification was performed using the PureYield™ Plasmid Miniprep System (column method, Promega). Plasmid concentration and purity were checked on a Nanodrop. Linear templates for PURE system reactions were prepared by PCR using the plasmids as substrates with primers 194 and 709 (**Supplementary Table 1**). Amplification products were checked on a 1% agarose gel and were purified using the Wizard SV Gel kit. DNA concentration and purity were measured using a ND-1000 UV-Vis Spectrophotometer (Nanodrop Technologies).

The DNA fragment containing the *zapA* gene (original sequence from *E. coli* K12 strain) was inserted in a pIDTSMART-AMP plasmid (Integrated DNA Technologies). The plasmid was transformed into *E. coli* TOP10 cells. Transformation, plasmid purification and production of linear DNA templates were performed as described above.

The *ftsA*_opt_ and *zapA*_opt_ constructs (starting with a T7 promoter and ending with the T7 terminator) were sequence-optimized for codon usage, GC content and 5’ mRNA secondary structures, and were inserted in a pJET1 and pUC57 plasmid, respectively (GeneScript). Plasmids were amplified and purified as described above. All sequences of the linearized constructs can be found in the Supplementary Methods.

### Purified proteins

Purified proteins were prepared and labelled according to published protocols^47^. FtsZ-Alexa647 (45 µM stock) was stored in a buffer containing 50 mM Tris, 500 mM KCl, 5 mM MgCl_2_ and 5% glycerol at pH 7.4. sZipA-Alexa488 (14.33 µM stock) was stored in a buffer containing 50 mM Tris, 50 mM KCl, and 1 mM EDTA at pH 7.4.

### Cell-free gene expression

PURE*frex*2.0 (GeneFrontier Corporation, Japan) was utilized following storing and handling instructions provided by the supplier. Linear DNA templates were used in single-gene expression assays at a final concentration of 5 nM. In co-expression experiments, both *ftsA*_opt_ and *zapA*_opt_ constructs were included at 5 nM and 10 nM, respectively, along with 1 µL of DnaK Mix (GeneFrontier Corporation). DnaK Mix consists of highly purified *E. coli* DnaK, DnaJ and GrpE chaperone proteins. Reactions of 20 µL volume were carried out in test tubes for 3 h at 37 °C. When indicated, samples were supplemented with purified proteins (FtsZ-Alexa647, sZipA-Alexa488) and added either on top of an SLB or used for lipid film swelling.

### Co-translational labelling of *in vitro* synthesized proteins and gel analysis

PURE*frex*2.0 reaction mixtures were supplemented with 0.5 μL of GreenLys (FluoroTect™ GreenLys, Promega) and gene expression was performed in a test tube as described above. Samples were treated with RNase (RNaseA Solution, Promega) for 30 min and proteins were denatured for 10 min at 90 °C in 2× SDS loading buffer with 10 mM DTT. Samples were loaded on a 18% SDS-PAGE (polyacrylamide gel electrophoresis) gel. Visualization of the fluorescently labelled translation products was performed on a fluorescence gel imager (Typhoon, Amersham Biosciences) using a 488-nm laser and a band pass emission filter of 520 nm.

### Fabrication and cleaning of the imaging chambers

Home-made glass chambers were used in both SLB and liposome experiments. Three microscopy glass slides (1-mm thick) were glued on top of each other with NOA 61 glue (Norland Products) and holes with a diameter of 2.5 mm were drilled. A 150 µm-thick coverslip (Menzel-Gläser, Germany) was glued with NOA 61 to cover the apertures, creating the bottom of glass chambers. Cleaning was performed by successive washing steps of 10 min each in a bath sonicator (Sonorex digitec, Bandelin), as follows: chloroform and methanol (1:1 volume ratio), 2% Hellmanex, 1 M KOH, 100% ethanol and finally MilliQ water. For SLB experiments the glass chambers were further treated every two to three experiments with Acid Piranha.

### Lipids

1,2-dioleoyl-sn-glycero-3-phosphocholine (DOPC), 1,2-dioleoyl-sn-glycero-3-phosphoethanolamine (DOPE), 1,2-dioleoyl-sn-glycero-3phosphoglycerol (DOPG), 1’,3’-bis[1,2-dioleoyl-sn-glycero-3-phospho]-glycerol (18:1 CL), 1,2-distearoyl-sn-glycero-3-phosphoethanolamine-N-[biotinyl(polyethylene glycol)-2000 (DSPE-PEG-biotin), and 1,2-dioleoyl-sn-glycero-3-[(N-(5-amino-1-carboxypentyl)iminodiacetic acid)succinyl] (DGS-NTA) were from Avanti Polar Lipids. Texas Red 1,2-dihexadecanoyl-sn-glycero-3-phosphoethanolamine (DHPE-TexasRed) was from Invitrogen.

### Preparation of small unilamellar vesicles

Small unilamellar vesicles (SUVs) were used as precursors for the formation of SLBs. Lipids DOPC (4 µmol), DOPG (1 µmol) and DGS-NTA (0.25 µmol), all dissolved in chloroform (Avanti Polar Lipids), were mixed in a glass vial. A lipid film was deposited on the wall of the vial upon solvent evaporation by applying a gentle flow of argon and was further desiccated for 30 min at room temperature. The lipid film was resuspended with 400 µL of SLB buffer (50 mM Tris, 300 mM KCl, 5 mM MgCl_2_, pH 7.5) and the solution was vortexed for a few minutes. The final lipid concentration was 1.25 mg mL^−1^. A two-step extrusion (each of 11 passages) was carried out using the Avanti mini extruder (Avanti Polar Lipids) equipped with 250 µL Hamilton syringes (Avant Polar Lipids), filters (drain disc 10 mm diameter, Whatman) and a polycarbonate membrane with a pore size of 0.2 µm (step 1) or 0.03 µm (step 2) (Nuclepore track-etched membrane, Whatman).

### Formation of supported lipid bilayers

The imaging chamber was treated with oxygen plasma (Harrick Plasma basic plasma cleaner) for 30 min to activate the glass surface. Immediately after plasma cleaning the SUV solution was added to the sample reservoir at a final lipid concentration of 0.94 mg mL^−1^ together with 3 mM CaCl_2_. The chamber was closed by sticking a coverslip using a double-sided adhesive silicone sheet (Life Technologies) and the sample was incubated for 30 min at 37 °C. Next, the chamber was opened and the SLB was carefully washed six times with SLB buffer. Under these conditions, the SLB contains 4.8 molar % of 18:1 DGS-NTA (Ni^2+^) lipids, which is within the range studied by ref.^27^ (0.5 – 10 mol%), similar as in ref.^28^ (1-8 mol%) but higher than in ref.^37^ (0.02 – 0.08 mol% of full length ZipA, DGS-NTA lipid was not used in this study) and lower than in ref.^47^ (10 mol%).

### Activity assays on supported membranes

In the experiments involving sZipA-Alexa488, 1 µM of the purified protein was first incubated on top of an SLB for 10 min at room temperature. The SLB was washed with 10 µL reaction buffer (50 mM Tris-HCl, 150 mM KCl, 5 mM MgCl_2_, pH 7.5). Then, 20 µL of sample (composition is specified where relevant) was added on top of the SLB and the chamber was sealed by sticking a 20×20 mm coverslip with a double-sided adhesive silicone sheet. In the experiments with FtsA, the *ftsA* or *ftsA*_opt_ gene was either directly expressed on top of the SLB, or in a test tube and subsequently added onto an SLB as part of the sample. In the earlier configuration, a 20 µL PURE*frex*2.0 reaction was carried out on top of an SLB and 10 µL were removed and replaced by the activity assay mixture. The exact composition of the sample varies for the different experiments and is specified where appropriate. In all cases, samples contained 2 mM GTP, supplemented with 2 mM ATP in FtsA experiments. In all assays without ZapA, Ficoll70 was added to a final concentration of 50 g L^−1^. No oxygen-scavenging system was used, unlike in ref.^28^ but like in ref.^37^.

### Spinning disk microscopy

Supported lipid bilayers were imaged with an Olympus iX81 inverted fluorescence microscope equipped with a 100 × oil immersion objective (Olympus), an iXon3 EMCCD camera (Andor Technology) and a Nipkow spinning disk (CSU-XI, Yokogawa). FtsZ-Alexa647 and sZipA-Alexa488 were imaged using a 640 nm and 491 nm laser line, respectively, and appropriate emission filters (685/40 nm or 525/50 nm). The software Andor IQ3 (Andor Technology Ltd.) was used for image acquisition and identical settings were used for all experiments. Experiments were conducted at room temperature.

### Preparation of lipid-coated beads

Lipid-coated beads were prepared according to our published protocol^35^ with the following lipid composition: DOPC (50 mol%), DOPE (36 mol%), DOPG (12 mol%), 18:1 CL (2 mol%), DSPE-PEG-biotin (1 mass%) and DHPE-TexasRed (0.5 mass%) for a total mass of 2 mg. Lipids dissolved in chloroform were mixed in a round-bottom glass flask. Methanol containing 100 mM rhamnose (Sigma-Aldrich) was added to the solution in a chloroform-to-methanol volume ratio of 2.5:1. Then, 1.5 g of 212-300 µm glass beads (acid washed, Sigma Aldrich) were poured to the lipid-rhamnose mixture and the organic solvent was removed by rotary evaporation at 200 mbar for 2 h at room temperature, followed by overnight desiccation. Lipid-coated beads were stored under argon at –20 °C until use.

### Production and immobilization of gene-expressing liposomes

A PURE*frex*2.0 reaction mixture was assembled as described above. Either or both *ftsA*_opt_ and *zapA*_opt_ DNA constructs were added at a final concentration of 5 nM and 10 nM, respectively. The solution was supplemented with (final concentrations indicated): 2 mM GTP, 2 mM ATP, 3 µM FtsZ-Alexa647 and MilliQ to reach a final volume of 20 µL. About 20 mg of lipid-coated was added to the solution and liposomes were formed by natural swelling of the lipid film for 2 h on ice, protected from light. During incubation, the tube was gently rotated manually a few times. Four freeze-thaw cycles were then applied by dipping the sample in liquid nitrogen and thawing on ice. The sample reservoir of the imaging chamber was functionalized with 1:1 molar ratio of bovine serum albumin (BSA) and BSA-biotin (1 mg mL^−1^, Thermo Fisher Scientific), and then with Neutravidin (1 mg mL^−1^, Sigma Aldrich), to tether the biotinylated liposomes. About 7 μL of the liposome solution was carefully pipetted (with a cut tip) into the imaging chamber and supplemented with RQ1 DNase (0.07 U µL^−1^) to preclude gene expression outside liposomes. The chamber was sealed by sticking a 20×20 mm coverslip with a double-sided adhesive silicone sheet. Expression was performed directly on the confocal microscope at 37 °C for 1.5 to 6 h.

### Confocal microscopy

A Nikon A1R Laser scanning confocal microscope equipped with an SR Apo TIRF 100 × oil immersion objective was used to image liposomes. The 561 nm and 640 nm laser lines were used in combination with appropriate emission filters to image the Texas Red membrane dye and FtsZ-Alexa647, respectively. The software NIS (Nikon) was used for image acquisition and identical settings were used for all experiments. Samples were mounted on a temperature-controlled stage maintained at 37 °C during imaging.

### Fluorescence recovery after photobleaching (FRAP)

FRAP experiments were performed on an Olympus iX81 spinning disk microscope. Images were acquired using the following protocol: 10 frames every s, 10 frames every 250 ms, 10 frames every 2 s, 10 frames every 4 s. Analysis of the FRAP images was performed with ImageJ^48,49^ using the *FRAP profiler* plug-in. The intensity of a bleached region of interest (ROI, 29×29 pixels) was measured over time and normalized to the intensity of the surrounding (250×250 pixels area centred on the ROI) to correct for the bleaching that occurs during image acquisition. Fitting of the FRAP curves was generated in GraphPad Software Inc. using a one-phase exponential model. At least three FRAP measurements were performed in each sample analysed.

### Structured illumination microscopy (SIM)

3D SIM images have been acquired with a Nikon SIM microscope equipped with a Nikon 100× and 1.49 NA Apo TIRF SR objective and a 640 nm laser line. The acquisition and reconstruction of the SIM images have been performed using the Nikon NIS element software. SIM RAW data and their corresponding reconstructed images were quality-checked using the Fiji plug-in *SIMcheck*^50^.

### Quantitative image analysis

Image analysis was performed using Mathematica (Wolfram Research, version 11.3). All images were corrected for uneven illumination by applying a Gaussian filter with radius 70 pixels to each image, fitting a third-degree polynomial to the filtered image, and dividing the original image pixel-by-pixel with the fitted polynomial. To segment the filaments a ridge filter with sigma=1 was applied and the resulting image binarized using morphological binarization with the default parameters. When needed this image was convolved with a Laplacian-of-Gaussian filter with radius 2 pixels and inversed (necessary for images with thin filaments). Filament thicknesses were calculated from this image as the distance of the centerline of filaments to the edge using the distance transform function. Branch point density and filament curvature were calculated after the thinning operation was applied to the segmented image. Curvatures were approximated at each pixel along the thinned filament with a distance larger than 2 pixels to the next branch point using Gaussian smoothing on the derivative functions. The image processing steps are illustrated in **Supplementary Fig. 3**.

## Acknowledgements

We thank Jeremie Capoulade and Duco Blanken for assistance with fluorescence microscopy, and Zohreh Nourian, Sanne Wiersma, Maryse Karsten and Mona Mohseni Kabir for preliminary experiments. Microscopy measurements were performed at the Kavli Nanolab Imaging Center Delft. This work was financially supported by the Netherlands Organization for Scientific Research (NWO/OCW) through the ‘BaSyC – Building a Synthetic Cell’ Gravitation grant (024.003.019) and the FOM programme nr.151, and by the Spanish government grant BFU2016-75471-C2-1-P.

## Author contribution

CD conceived and supervised the research. EG, JN and CD designed the experiments. EG, JN and IZ performed the experiments. EG, JN, IZ, AD and CD analyzed the data. CD and EG wrote the paper. MJ and GR provided the purified FtsZ and sZipA proteins. All authors discussed the results and gave inputs on the manuscript.

## Supplementary information

The Supplementary Information is available on the journal website. It includes Supplementary Methods, Supplementary Notes, as well as Supplementary Tables, Figures and a Movie.

## Conflict of interest

The authors declare no conflict of interest.

## REFERENCES

1. Shimizu, Y. et al. Cell-free translation reconstituted with purified components. Nat. Biotechnol. 19, 751–755 (2001).

2. Gros, J., Devbhandari, S. & Remus, D. Origin plasticity during budding yeast DNA replication in vitro. EMBO J. 33, 621–636 (2014).

3. Lee, K., Gallop, J. L., Rambani, K. & Kirschner, M. W. Self-Assembly of filopodia-like structures on supported lipid bilayers. Science. 329, 1341–1345 (2010).

4. Vignaud, T., Blanchoin, L. & Théry, M. Directed cytoskeleton self-organization. Trends in Cell Biology vol. 22 671–682 (2012).

5. Nguyen, P. A. et al. Spatial organization of cytokinesis signaling reconstituted in a cell-free system. Science. 346, 244–247 (2014).

6. Garner, E. C., Campbell, C. S., Weibel, D. B. & Mullins, R. D. Reconstitution of DNA segregation driven by assembly of a prokaryotic actin homolog. Science. 315, 1270–1274 (2007).

7. Dannhauser, P. N. & Ungewickell, E. J. Reconstitution of clathrin-coated bud and vesicle formation with minimal components. Nat. Cell Biol. 14, 634–639 (2012).

8. Focus on the benefits of building life’s systems from scratch. Nature 563, 155–155 (2018).

9. Nourian, Z., Scott, A. & Danelon, C. Toward the assembly of a minimal divisome. Syst. Synth. Biol. 8, 237 (2014).

10. Scott, A. et al. Cell-free phospholipid biosynthesis by gene-encoded enzymes reconstituted in liposomes. PLoS One 11, e0163058 (2016).

11. van Nies, P. et al. Self-replication of DNA by its encoded proteins in liposome-based synthetic cells. Nat. Commun. 9, 1583 (2018).

12. Godino, E. et al. De novo synthesized Min proteins drive oscillatory liposome deformation and regulate FtsA-FtsZ cytoskeletal patterns. Nat. Commun. 10, 4969 (2019).

13. Bi, E. & Lutkenhaus, J. FtsZ ring structure associated with division in *Escherichia coli*. Nature 354, 161–164 (1991).

14. Ma, X., Ehrhardt, D. W. & Margolin, W. Colocalization of cell division proteins FtsZ and FtsA to cytoskeletal structures in living *Escherichia coli* cells by using green fluorescent protein. Proc. Natl. Acad. Sci. 93, 12998–13003 (1996).

15. Hale, C. A. & de Boer, P. A. Recruitment of ZipA to the septal ring of *Escherichia coli* is dependent on FtsZ and independent of FtsA. J. Bacteriol. 181, 167–76 (1999).

16. Walker, B. E., Männik, J. & Mannik, J. Transient membrane-linked FtsZ assemblies precede Z-ring formation in *Escherichia Coli*. Current Biology 30, 499–508 (2019).

17. de Boer, P., Crossley, R. & Rothfield, L. The essential bacterial cell-division protein FtsZ is a GTPase. Nature 359, 254–256 (1992).

18. Mukherjee, A. & Lutkenhaus, J. Guanine nucleotide-dependent assembly of FtsZ into filaments. J. Bacteriol. 176, 2754–8 (1994).

19. Pichoff, S. & Lutkenhaus, J. Unique and overlapping roles for ZipA and FtsA in septal ring assembly in *Escherichia coli*. EMBO J. 21, 685–93 (2002).

20. Pichoff, S. & Lutkenhaus, J. Tethering the Z ring to the membrane through a conserved membrane targeting sequence in FtsA. Mol. Microbiol. 55, 1722–1734 (2005).

21. Hale, C. A. & de Boer, P. A. Direct binding of FtsZ to ZipA, an essential component of the septal ring structure that mediates cell division in *E. coli*. Cell 88, 175–85 (1997).

22. Ortiz, C., Natale, P., Cueto, L. & Vicente, M. The keepers of the ring: regulators of FtsZ assembly. FEMS Microbiol. Rev. 40, 57–67 (2016).

23. Osawa, M. & Erickson, H. P. Liposome division by a simple bacterial division machinery. Proc. Natl. Acad. Sci. U. S. A. 110, 11000–4 (2013).

24. Cabré, E. J. et al. Bacterial division proteins FtsZ and ZipA induce vesicle shrinkage and cell membrane invagination. J. Biol. Chem. 288, 26625–34 (2013).

25. Ramirez-Diaz, D. A. et al. Treadmilling analysis reveals new insights into dynamic FtsZ ring architecture. PLOS Biol. 16, e2004845 (2018).

26. Krupka, M. et al. *Escherichia coli* FtsA forms lipid-bound minirings that antagonize lateral interactions between FtsZ protofilaments. Nat. Commun. 8, 15957 (2017).

27. Krupka, M., Sobrinos-Sanguino, M., Jiménez, M., Rivas, G. & Margolin, W. *Escherichia coli* ZipA organizes FtsZ Polymers into dynamic ring-like protofilament structures. MBio 9, e01008–18 (2018).

28. Loose, M. & Mitchison, T. J. The bacterial cell division proteins FtsA and FtsZ self-organize into dynamic cytoskeletal patterns. Nat. Cell Biol. 16, 38–46 (2014).

29. Osawa, M., Anderson, D. E. & Erickson, H. P. Curved FtsZ protofilaments generate bending forces on liposome membranes. EMBO J. 28, 3476–84 (2009).

30. Szwedziak, P., Wang, Q., Bharat, T. A. M., Tsim, M. & Löwe, J. Architecture of the ring formed by the tubulin homologue FtsZ in bacterial cell division. Elife 3, e04601 (2014).

31. Söderström, B. et al. Disassembly of the divisome in *Escherichia coli* : evidence that FtsZ dissociates before compartmentalization. Mol. Microbiol. 92, 1–9 (2014).

32. Daley, D. O., Skoglund, U. & Söderström, B. FtsZ does not initiate membrane constriction at the onset of division. Sci. Rep. 6, 33138 (2016).

33. Furusato, T. et al. *De novo* synthesis of basal bacterial cell division proteins FtsZ, FtsA, and ZipA inside giant vesicles. ACS Synth. Biol. 7, 953–961 (2018).

34. Doerr, A. et al. Modelling cell-free RNA and protein synthesis with minimal systems. Phys. Biol. 16, 025001 (2019).

35. Blanken, D., van Nies, P. & Danelon, C. Quantitative imaging of gene-expressing liposomes reveals rare favorable phenotypes. Phys. Biol. 16, 045002 (2019).

36. Caldas, P., López-Pelegrín, M., Pearce, D.J.G. et al. Cooperative ordering of treadmilling filaments in cytoskeletal networks of FtsZ and its crosslinker ZapA. Nat. Commun. 10, 5744 (2019).

37. Martos, A. et al. FtsZ polymers tethered to the membrane by ZipA are susceptible to spatial regulation by Min waves. Biophys. J. 108, 2371–2383 (2015).

38. Rivas, G., Alfonso, C., Jiménez, M., Monterroso, B. & Zorrilla, S. Macromolecular interactions of the bacterial division FtsZ protein: from quantitative biochemistry and crowding to reconstructing minimal divisomes in the test tube. Biophys. Rev. 5, 63–77 (2013).

39. Rueda, S., Vicente, M. & Mingorance, J. Concentration and assembly of the division ring proteins FtsZ, FtsA, and ZipA during the *Escherichia coli* cell cycle. J. Bacteriol. 185, 3344–51 (2003).

40. Osawa, M. & Erickson, H. P. Inside-out Z rings - constriction with and without GTP hydrolysis. Mol. Microbiol. 81, 571–579 (2011).

41. Geissler, B., Elraheb, D. & Margolin, W. A gain-of-function mutation in ftsA bypasses the requirement for the essential cell division gene zipA in *Escherichia coli*. Proc. Natl. Acad. Sci. 100, 4197–4202 (2003).

42. Pichoff, S., Shen, B., Sullivan, B. & Lutkenhaus, J. FtsA mutants impaired for self-interaction bypass ZipA suggesting a model in which FtsA’s self-interaction competes with its ability to recruit downstream division proteins. Mol. Microbiol. 83, 151–167 (2012).

43. Osawa, M., Anderson, D. E. & Erickson, H. P. Reconstitution of contractile FtsZ rings in liposomes. Science. 320, 792–794 (2008).

44. Cabré, E. J. et al. The nucleoid occlusion SlmA protein accelerates the disassembly of the FtsZ protein polymers without affecting their GTPase activity. PLoS One 10, e0126434 (2015).

45. Monterroso, B. et al. The bacterial DNA binding protein MatP involved in linking the nucleoid terminal domain to the divisome at midcell interacts with lipid membranes. MBio 10, e00376–19 (2019).

46. Noireaux, V., Bar-Ziv, R. & Libchaber, A. Principles of cell-free genetic circuit assembly. Proc. Natl. Acad. Sci. 100, 12672–12677 (2003).

47. Mateos-Gil, P. et al. FtsZ polymers bound to lipid bilayers through ZipA form dynamic two dimensional networks. Biochim. Biophys. Acta - Biomembr. 1818, 806–813 (2012).

48. Schindelin, J. et al. ImageJ2: ImageJ for the next generation of scientific image data. BMC Bioinformatics. 18, 529 (2017).

49. Schneider, C. A., Rasband, W. S., & Eliceiri, K. W. NIH Image to ImageJ: 25 years of image analysis. Nat. Methods. 9, 671–675 (2012).

50. Ball, G. et al. SIMcheck: a toolbox for successful super-resolution structured illumination microscopy. Sci. Rep. 5, 15915 (2015).

