## Supplementary Information for "Cell-free biogenesis of bacterial division proto-rings that can constrict liposomes"

**For**

### SUPPLEMENTARY METHODS

#### Sequence of the *ftsA* construct

TAATACGACTCACTATAGGGGAATTGTGAGCGGATAACAATTCCCCTCTAGAAATAATTTTGTTTAACTTTAAG
AAGGAGATATACATatgATCAAGGCGACGGACAGAAAACTGGTAGTAGGACTGGAGATTGGTACCGCGAAGG
TTGCCGCTTTAGTAGGGGAAGTTCTGCCCGACGGTATGGTCAATATCATTGGCGTGGGCAGCTGCCCGTCGCG
TGGTATGGATAAAGGCGGGGTGAACGACCTCGAATCCGTGGTCAAGTGCGTACAACGCGCCATTGACCAGGC
AGAATTGATGGCAGATTGTCAGATCTCTTCGGTATATCTGGCGCTTTCTGGTAAGCACATCAGCTGCCAGAAT
GAAATTGGTATGGTGCCTATTTCTGAAGAAGAAGTGACGCAAGAAGATGTGGAAAACGTCGTCCATACCGCG
AAATCGGTGCGTGTGCGCGATGAGCATCGTGTGCTGCATGTGATCCCGCAAGAGTATGCGATTGACTATCAG
GAAGGGATCAAGAATCCGGTAGGACTTTCGGGCGTGCGGATGCAGGCAAAAGTGCACCTGATCACATGTCAC
AACGATATGGCGAAAAACATCGTCAAAGCGGTTGAACGTTGTGGGCTGAAAGTTGACCAACTGATATTTGCC
GGACTGGCATCAAGTTATTCGGTATTGACGGAAGATGAACGTGAACTGGGTGTCTGCGTCGTCGATATCGGT
GGTGGTACAATGGATATCGCCGTTTATACCGGTGGGGCATTGCGCCACACTAAGGTAATTCCTTATGCTGGCA
ATGTCGTGACCAGTGATATCGCTTACGCCTTTGGCACGCCGCCAAGCGACGCCGAAGCGATTAAAGTTCGCCA
CGGTTGTGCGCTGGGTTCCATCGTTGGAAAAGATGAGAGCGTGGAAGTGCCGAGCGTAGGTGGTCGTCCGC
CACGGAGTCTGCAACGTCAGACACTGGCAGAGGTGATCGAGCCGCGCTATACCGAGCTGCTCAACCTGGTCA
ACGAAGAGATATTGCAGTTGCAGGAAAAGCTTCGCCAACAAGGGGTTAAACATCACCTGGCGGCAGGCATTG
TATTAACCGGTGGCGCAGCGCAGATCGAAGGTCTTGCAGCCTGTGCTCAGCGCGTGTTTCATACGCAAGTGC
GTATCGGCGCGCCGCTGAACATTACCGGTTTAACGGATTATGCTCAGGAGCCGTATTATTCGACGGCGGTGG
GATTGCTTCACTATGGGAAAGAGTCACATCTTAACGGTGAAGCTGAAGTAGAAAAACGTGTTACAGCATCAGT
TGGCTCGTGGATCAAGCGACTCAATAGTTGGCTGCGAAAAGAGTTTtaaGGATCCGGCTGCTAACAAAGCCCG
AAAGGAAGCTGAGTTGGCTGCTGCCACCGCTGAGCAATAACTAGCATAACCCCTTGGGGCCTCTAAACGGGT
CTTGAGGGGTTTTTTG

**Sequence of the optimized *ftsA*$_{opt}$ construct**

TAATACGACTCACTATAGGGGAATTGTGAGCGGATAACAATTCCCCTCTAGAAATAATTTTGTTTAACTTTAAG
AAGGAGATATACATATGATCAAGGCGACCGACCGTAAGCTGGTTGTGGGCCTGGAGATTGGCACCGCGAAG
GTTGCGGCGCTGGTTGGCGAGGTTCTGCCGGATGGTATGGTTAACATTATCGGCGTTGGTAGCTGCCCGAGC
CGTGGCATGGACAAAGGTGGTGTGAACGACCTGGAAAGCGTGGTTAAGTGCGTGCAGCGTGCGATTGACCA
GGCGGAGCTGATGGCGGACTGCCAAATCAGCAGCGTTTACCTGGCGCTGAGCGGCAAGCACATCAGCTGCCA
AAACGAGATTGGTATGGTGCCGATTAGCGAAGAGGAAGTTACCCAGGAAGATGTGGAGAACGTGGTTCACA
CCGCGAAAAGCGTTCGTGTGCGTGATGAACACCGTGTGCTGCACGTTATCCCGCAAGAATACGCGATCGATTA
CCAGGAAGGTATCAAAAACCCGGTTGGTCTGAGCGGTGTTCGTATGCAGGCGAAAGTGCACCTGATTACCTG
CCACAACGATATGGCGAAGAACATTGTGAAAGCGGTTGAACGTTGCGGTCTGAAGGTTGACCAGCTGATCTT
CGCGGGTCTGGCGAGCAGCTACAGCGTTCTGACCGAAGATGAGCGTGAACTGGGTGTTTGCGTTGTGGATAT
CGGCGGTGGCACGATGGATATCGCGGTGTATACCGGTGGCGCGCTGCGTCACACCAAAGTGATTCCGTATGC
GGGTAACGTGGTTACCAGCGACATCGCGTACGCGTTTGGCACCCCGCCGAGCGATGCGGAGGCGATCAAAGT
GCGTCACGGTTGCGCGCTGGGTAGCATTGTGGGTAAAGATGAGAGCGTGGAAGTTCCGAGCGTTGGTGGCC
GTCCGCCGCGTAGCCTGCAACGTCAGACCCTGGCGGAAGTTATCGAGCCGCGTTACACCGAACTGCTGAACCT
GGTGAACGAAGAGATCCTGCAACTGCAAGAGAAACTGCGTCAGCAAGGTGTTAAGCACCACCTGGCGGCGG
GCATTGTTCTGACCGGCGGTGCGGCGCAGATCGAAGGTCTGGCGGCGTGCGCGCAACGTGTTTTCCACACCC
AAGTTCGTATCGGTGCGCCGCTGAACATCACCGGTCTGACCGATTACGCGCAAGAGCCGTACTATAGCACCGC
GGTTGGTCTGCTGCACTATGGCAAAGAGAGCCACCTGAACGGCGAGGCGGAAGTGGAGAAGCGTGTGACCG
CGAGCGTTGGTAGCTGGATTAAGCGTCTGAATAGCTGGCTGCGTAAGGAGTTCTAAGGATCCGGCTGCTAAC
AAAGCCCGAAAGGAAGCTGAGTTGGCTGCTGCCACCGCTGAGCAATAACTAGCATAACCCCTTGGGGCCTCT
AAACGGGTCTTGAGGGGTTTTTTG

**Sequence of the *zapA* construct**

TAATACGACTCACTATAGGGGAATTGTGAGCGGATAACAATTCCCCTCTAGAAATAATTTTGTTTAACTTTAAG
AAGGAGATATACATATGTCTGCACAACCCGTCGATATCCAAATTTTTGGCCGTTCACTGCGTGTGAACTGCCCG
CCTGACCAAAGGGATGCGTTGAATCAGGCAGCGGACGATCTGAACCAACGGTTGCAAGATCTGAAAGAACGC
ACTAGAGTCACAAATACTGAACAGTTGGTCTTCATTGCCGCATTGAATATCAGCTATGAGTTAGCGCAAGAAA
AAGCAAAGACTCGTGACTACGCGGCAAGTATGGAACAGCGTATTCGGATGCTGCAGCAGACCATAGAACAAG
CGTTACTTGAACAAGGTCGCATCACCGAAAAAACTAACCAAAACTTTGAATGAGGATCCGGCTGCTAACAAAG
CCCGAAAGGAAGCTGAGTTGGCTGCTGCCACCGCTGAGCAATAACTAGCATAACCCCTTGGGGCCTCTAAAC
GGGTCTTGAGGGGTTTTTTG

**Sequence of the optimized *zapA*$_{opt}$ construct**

TAATACGACTCACTATAGGGGAATTGTGAGCGGATAACAATTCCCCTCTAGAAATAATTTTGTTTAACTTTAAG
AAGGAGATATACATATGAGCGCGCAACCGGTGGACATCCAGATTTTTGGCCGTAGCCTGCGTGTGAACTGCC
CGCCGGACCAACGTGATGCGCTGAACCAGGCGGCGGACGATCTGAACCAGCGTCTGCAAGACCTGAAGGAG
CGTACCCGTGTGACCAACACCGAACAGCTGGTTTTCATCGCGGCGCTGAACATTAGCTACGAGCTGGCGCAG
GAAAAGGCGAAAACCCGTGATTATGCGGCGAGCATGGAGCAACGTATCCGTATGCTGCAACAAACCATTGAA
CAGGCGCTGCTGGAACAGGGTCGCATCACCGAGAAGACCAATCAGAATTTTGAATAAGGATCCGGCTGCTAA
CAAAGCCCGAAAGGAAGCTGAGTTGGCTGCTGCCACCGCTGAGCAATAACTAGCATAACCCCTTGGGGCCTC
TAAACGGGTCTTGAGGGGTTTTTTG

### SUPPLEMENTARY NOTES

#### Supplementary Note 1: Bundling of FtsZ filaments is required for the formation of long ZipA-FtsZ cytoskeletal structures in low-volume SLB assays

We examined the ability of purified FtsZ and the cytoplasmic domain of ZipA, sZipA[1], to assemble into membrane-tethered protofilaments in a PURE system background. Specifically, we attempted to reproduce previous results using purified FtsZ and sZipA in SLB assays[2]. To simultaneously visualize both proteins, FtsZ and sZipA were labelled with Alexa Fluor-647 and 488, respectively. Membrane recruitment and spatial organization of the proteins were imaged using spinning disk fluorescence microscopy. We used 3 µM FtsZ-A647, which is above the critical concentration for in vitro polymerization[3,4], and about 1 µM sZipA-A488 in the presence of GTP. The 4.8 mol% of DOGS lipid results in a regime of high-density of sZipA. While sZipA-conditional recruitment of FtsZ to the membrane was confirmed, no visible large-scale filaments could develop under these conditions, in contrast with previous observations[2]. To our surprise, repeating the experiment in the same minimal buffer as in Loose et al.[2] in place of PURE*frex*2.0 failed to reproduce filament networks (**Fig. 2a,b**). Given our unconventional setup for SLB assays which is designed for low-volume PURE system solutions, we asked whether the chamber dimension and geometry could cause this different behaviour. In both PURE*frex*2.0 background and minimal buffer, supplementing the solution with 12.5% Ficoll70, an artificial crowding agent known to promote FtsZ filament condensation[5], triggered the formation of a dense network of sZipA-FtsZ co-filaments (**Fig. 2 c,d**, **Supplementary Fig. 5**) having a similar morphology as previously reported[2]. The presence of long (>10 µm) bundles is typical to high membrane coverage with sZipA. At low-density sZipA and identical concentration of Ficoll70, only short bundles and three-dimensional networks of FtsZ were observed[6]. Although different areas of an SLB can show various protein network phenotypes, more condensed bundles were observed in PURE*frex*2.0 compared to the minimal buffer (**Fig. 2d**, **Supplementary Fig. 5**). Once formed, these structures are stable in time (at least up to 3 hours) with no large-scale reorganization within minutes. Fluorescence recovery after photobleaching (FRAP) experiments revealed that membrane-bound sZipA is immobile within the cytoskeletal network (**Supplementary Fig. 10**).

Collectively, these results indicate that the Ficoll70-induced sZipA-FtsZ filament networks have a different morphology in the physiological PURE*frex*2.0 background compared to the minimal buffer. Moreover, the total reaction volume acts as a control parameter in the formation of cytoskeletal patterns in SLB assays.

The above experiments involved Ficoll70 to promote lateral interaction of FtsZ filaments[6–9]. Alternatively, ZapA was produced in PURE*frex*2.0 starting from its native gene sequence and its ability to crosslink FtsZ filaments was examined using purified FtsZ on sZipA-membranes (**Fig. 2e,f**). Protein networks of typical flower-like morphology with less condensed filaments than in the presence of Ficoll70 were observed after 3 h expression (**Fig. 2f**). We noticed that these filament structures are not stable in time and vanished after a few seconds imaging (**Supplementary Fig. 6**). Because gene sequence optimization resulted in a higher yield of synthesized ZapA (**Supplementary Fig. 9**), this engineered construct was also tested in the activity assay. Expression of the *zapA*$_{opt}$ DNA template triggered the formation of a dense network of sZipA-FtsZ co-filaments that are stable in time (**Supplementary Fig. 7**). We attribute these new properties to the higher concentration of ZapA when expressed from the *zapA*$_{opt}$ gene (**Supplementary Fig. 9**). Different areas of the SLB show various protein network phenotypes ranging from flower-like morphology to long condensed bundles (**Supplementary Fig. 7**). Although many filaments were localizing on the membrane, sZipA-FtsZ co-filaments were abundantly found to extend in the bulk phase (**Supplementary Fig. 7)**. This suggests that the pre-assembly of ZapA-FtsZ bundles compromises the ability of sZipA-FtsZ to stably bind the membrane. We verified that cell-free synthesized ZapA is unable to produce membrane-bound FtsZ filaments in the absence of ZipA (**Supplementary Fig. 8**).

The exchange dynamics of FtsZ and sZipA subunits within polymers were quantified by FRAP. With Ficoll70, no recovery of sZipA signal was observed indicating poor lateral diffusion and turnover of the membrane-bound sZipA (**Supplementary Fig. 10**). In contrast, ZapA-mediated bundling of FtsZ filaments does not impair recovery of sZipA signal (**Supplementary Fig. 10**). Furthermore, FRAP experiments show that FtsZ-A647 monomers undergo twice faster exchange rate between filaments and bulk with expressed ZapA than with Ficoll70 (**Supplementary Fig. 10**).

All together these results underline the importance of using ZapA to drive physiological filament network morphology and subunit turnover in membrane-bound sZipA-FtsZ cytoskeletal structures. Furthermore, the experiments highlight that ZapA concentration is important for long-term stability of filaments.

### SUPPLEMENTARY MOVIE

**Supplementary Movie 1:** Time series of confocal fluorescence images showing that in-liposome synthesized FtsA assembles with FtsZ into ring-like structures that drive membrane neck formation and vesicle budding. Ring-forming protein clusters localized at a constriction site can split, which induces multiple necks separated by blebbing vesicles. The $ftsA_{\mathrm{opt}}$ DNA template was expressed within phospholipid vesicles in the presence of 3 µM purified FtsZ-A647. Green signal, membrane dye fluorescence; magenta signal, FtsZ-A647 fluorescence. Scale bar represents 10 µm.

### SUPPLEMENTARY TABLE

**Supplementary Table 1:** List of primers used in this study.

| Name | Sequence (5′ → 3′) | Comment |
|---|---|---|
| 709 | CAAAAAACCCCTCAAGACCCGTTTAGAGG | Anneals at the T7 terminator |
| 757 | TAATACGACTCACTATAGGG | Anneals at the T7 promoter |
| 376 | CTTTCGGGCTTTGTTAGCAGCCGGATCCTTAAAACTCTTTTCGCAGCCAAC | *ftsA* (with overhang for pET11-a) |
| 508 | TTTGTTTAACTTTAAGAAGGAGATATACATATGATCAAGGCGACGGACAG | *ftsA* (with overhang for pET11-a) |
| 194 | TAATACGACTCACTATAGGGGAATTGTGAGCGGATAACAATTCCCCT | Anneals at the T7 promoter |

### SUPPLEMENTARY FIGURES

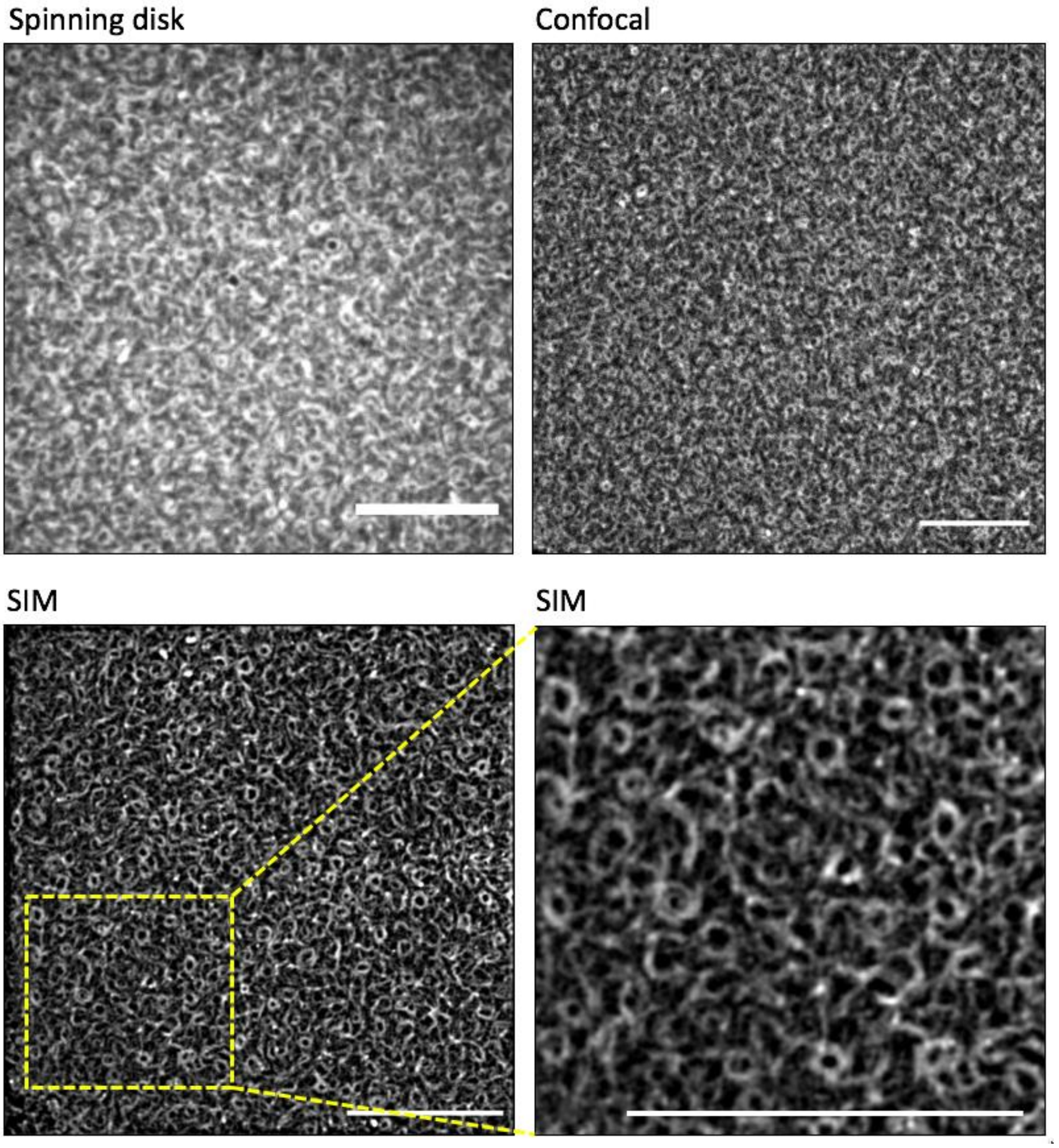


**Supplementary Fig. 1: Cell-free synthesized FtsA shapes FtsZ filaments into characteristic ring-like structures.** In situ expressed FtsA is sufficient to promote assembly of short and curved filaments of FtsZ (3 µM of purified FtsZ-A647) in the absence of Ficoll70. The sequence-optimized *ftsA*$_{opt}$ gene was used. Sample preparation was the same as in the main text **Fig. 1a,b**. Different fluorescence imaging modes have been employed: spinning disk, laser scanning confocal and structured illumination microscopy (SIM). Scale bars indicate 10 µm. SIM provides a higher spatial resolution, while spinning disk offers faster image acquisition.

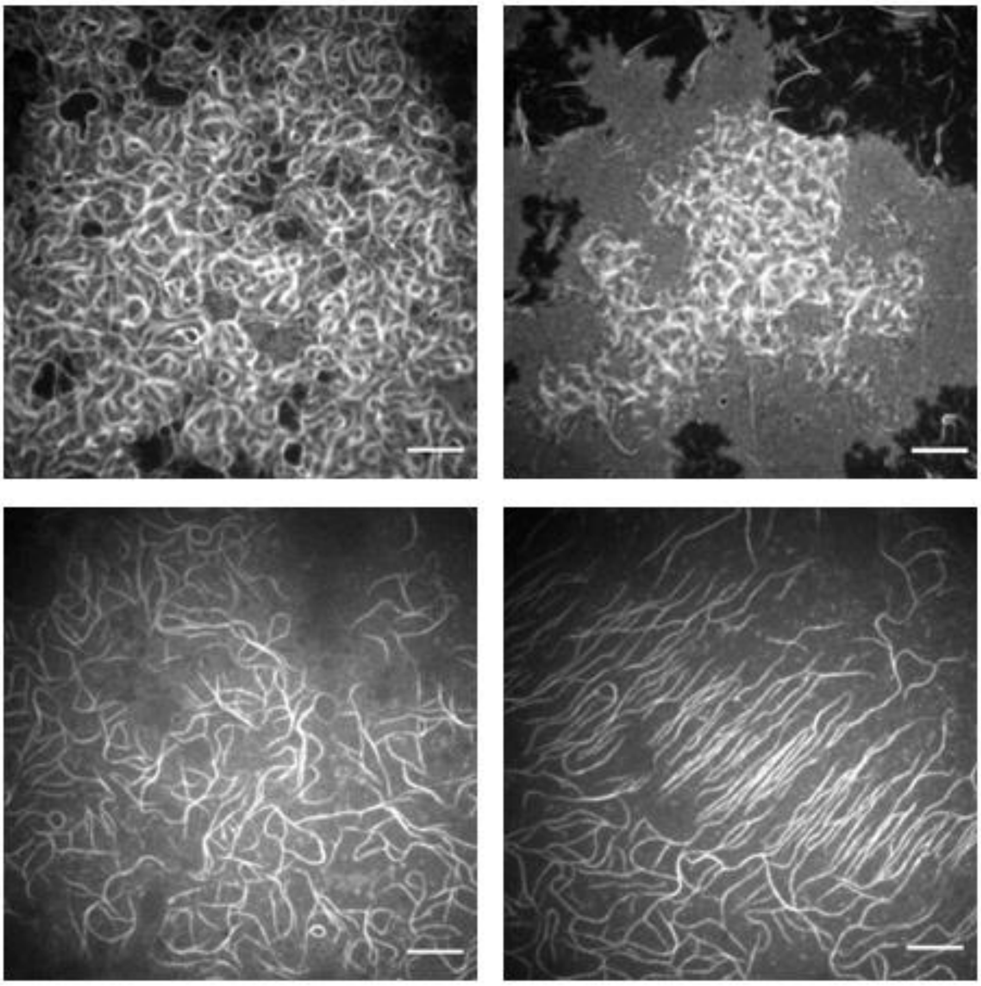

**Supplementary Fig. 2: Cytoskeletal filaments of FtsZ in the presence of co-expressed FtsA and ZapA proteins on an SLB.** FtsA and ZapA were directly expressed from their sequence-optimized constructs on an SLB, which drives the formation of thick FtsZ (3 µM of purified FtsZ-A647) filaments developing in large-scale networks with various morphologies. Four different fields of view are shown. Images were acquired in the FtsZ-A647 channel. Experimental conditions are similar as in the main text **Fig. 1g,h**. Scale bars indicate 10 µm.

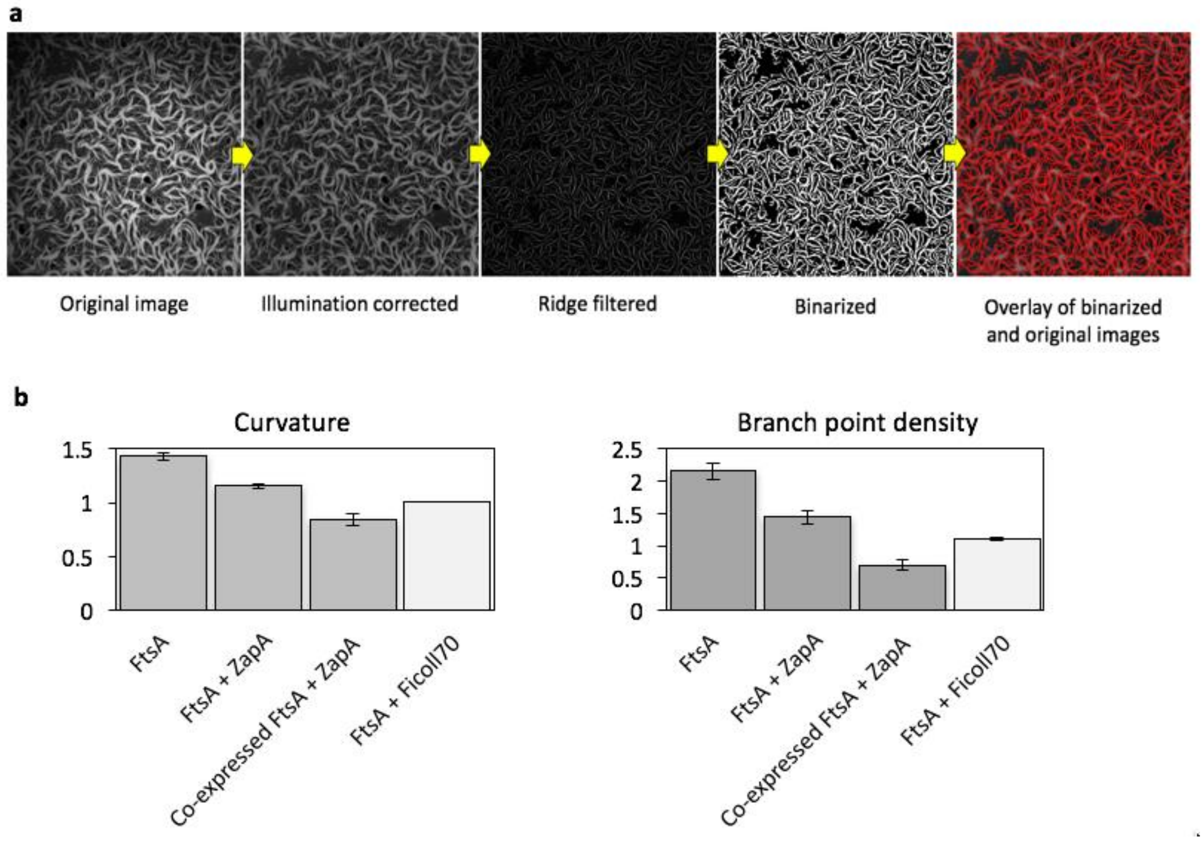


**Supplementary Fig. 3: Quantitative image analysis of cytoskeletal network properties. a**, Image processing workflow. **b**, Extracted parameters from the binarized images include filament curvature and branch point density. Four different fields of view have been analysed per experimental condition. Mean ± standard deviation values are reported. Curvature, $\mu m^{-1}$; branch point density, $\mu m^{-2}$.

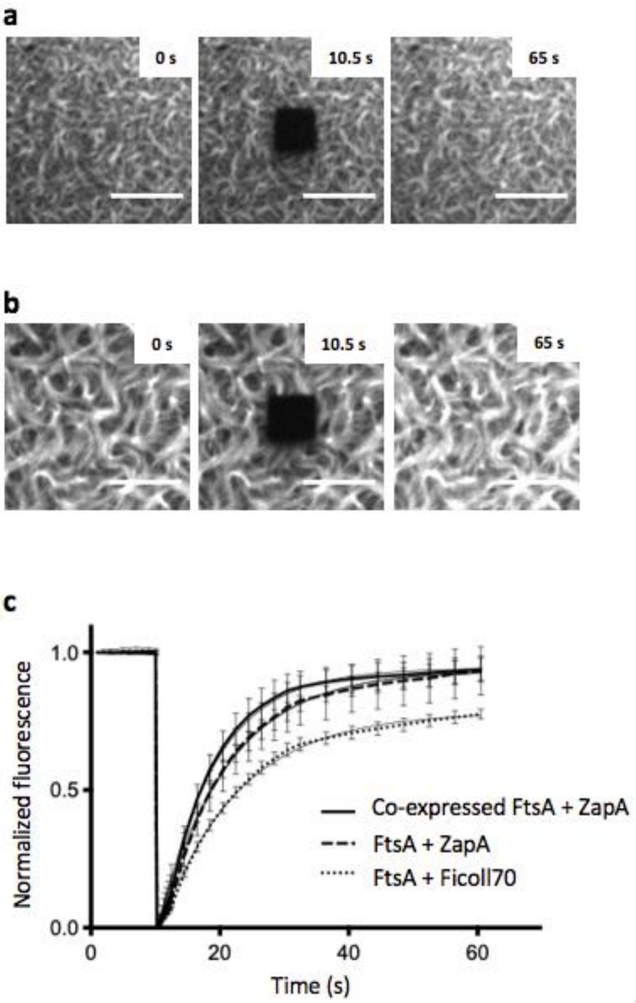


**Supplementary Fig. 4: FRAP experiments with purified FtsZ-A647 and cell-free expressed FtsA.** **a**, Time-lapsed fluorescence images of 3 µM FtsZ-A647 and cell-free expressed FtsA on an SLB in the presence of Ficoll70. The optimized $ftsA_{opt}$ construct was used. Scale bars represent 10 µm. **b**, Time-lapsed fluorescence images of 3 µM FtsZ-A647 and separately expressed FtsA and ZapA on an SLB. The optimized $ftsA_{opt}$ and native *zapA* constructs were used. Scale bars represent 10 µm. **c**, FRAP curves showing FtsZ monomer turnover under different experimental conditions, as annotated. Data are mean ± standard deviation values from four FRAP measurements per condition. Mono-exponential fits are appended. When bundling of FtsZ filaments was promoted by ZapA, a recovery halftime value of ~7 ± 2 s was found with both separately and co-expressed FtsA. With Ficoll70, the recovery halftime value is 8.4 ± 1.9 s, which is markedly lower than with sZipA (17.6 ± 1.3 s, **Supplementary Fig. 10**). These values are also similar (7.5 ± 2.4 s) for FtsZ turnover in protofilaments seeded on sZipA membranes with ZapA (**Supplementary Fig. 10**).

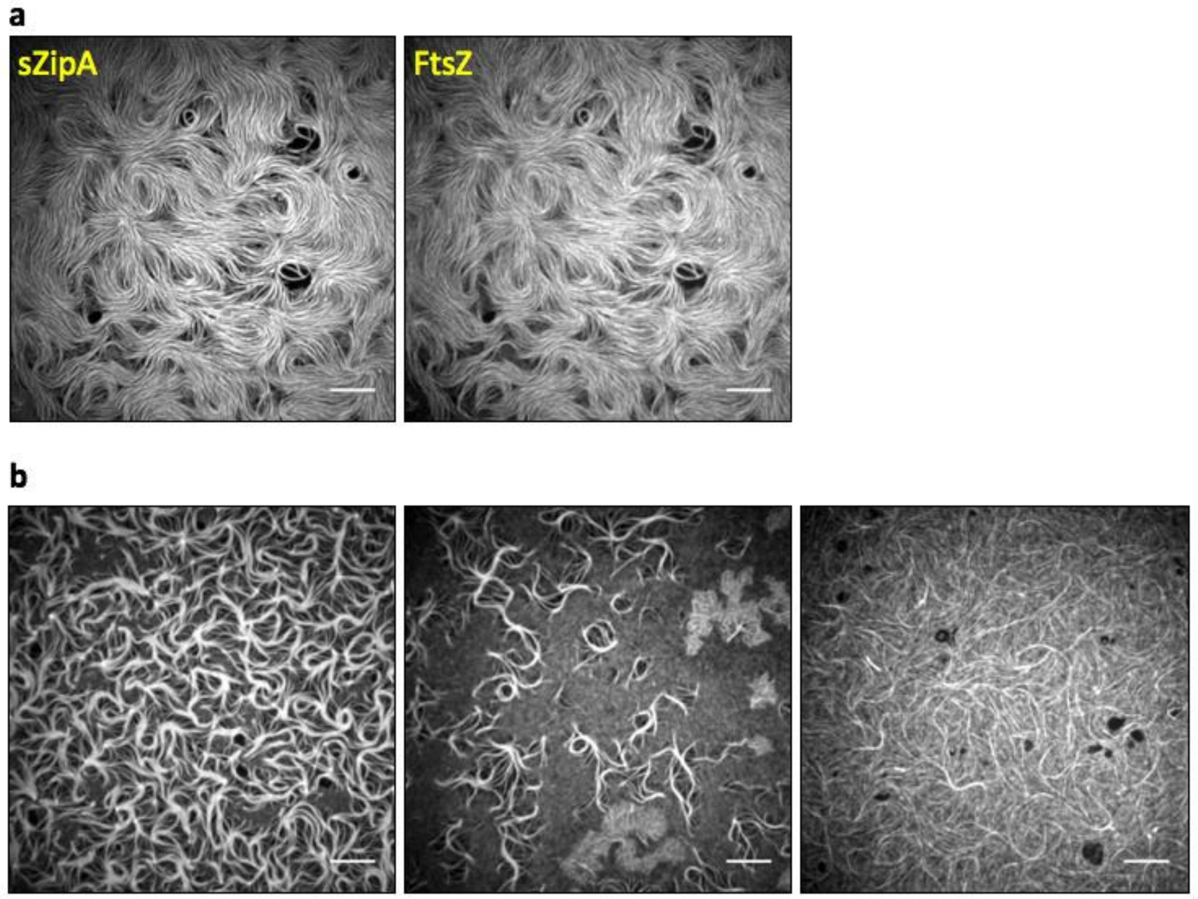


**Supplementary Fig. 5: Heterogeneity of FtsZ polymer network morphologies on sZipA-bound membranes. a**, Fluorescence images of sZipA-A488 (left) and FtsZ-A647 (right) in the minimal reaction buffer with Ficoll70. In the absence of Ficoll70, no large-scale filament networks develop (see main text **Fig. 2a,b**). **b**, Images of purified FtsZ-647 (3 µM) and sZipA-488 (1 µM) on top of an SLB in PURE*frex*2.0 background supplemented with Ficoll70 (12.5% m/v). Three typical fields of view showing different protein network morphologies are reported. Images in the FtsZ-A647 channel are shown. Scale bars represent 10 µm.

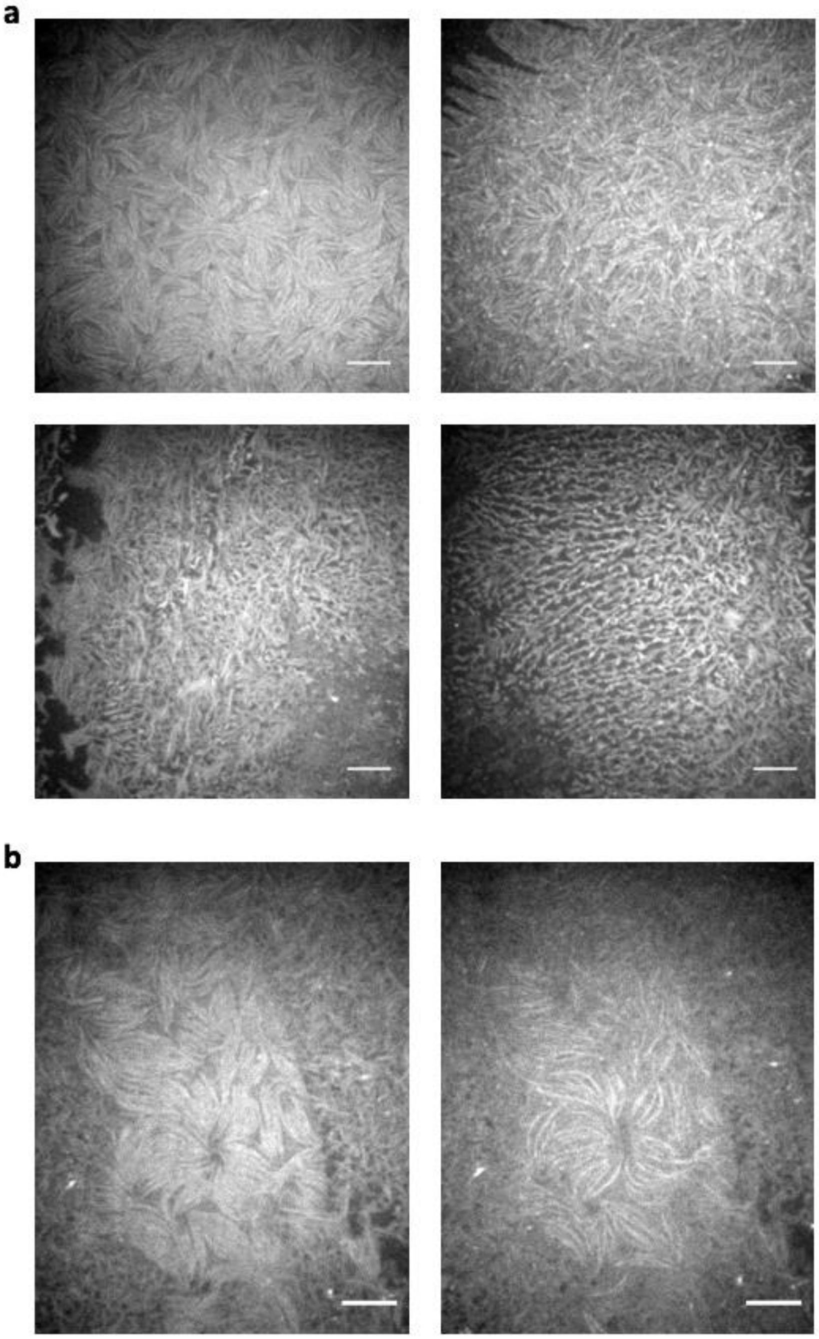


**Supplementary Fig. 6: Instability and morphological heterogeneity of ZapA-mediated FtsZ polymer networks. a**, ZapA, cell-free synthesized from the native gene sequence, promotes formation of FtsZ bundles (3 µM of purified FtsZ-A647) recruited on an SLB pre-incubated with 1 µM purified sZipA-A488. Four different fields of view in the FtsZ-A647 channel are shown. **b**, Time-lapsed images of the same SLB region showing disassembly of FtsZ bundles over a few seconds imaging (from left to right). Scale bars indicate 10 µm.

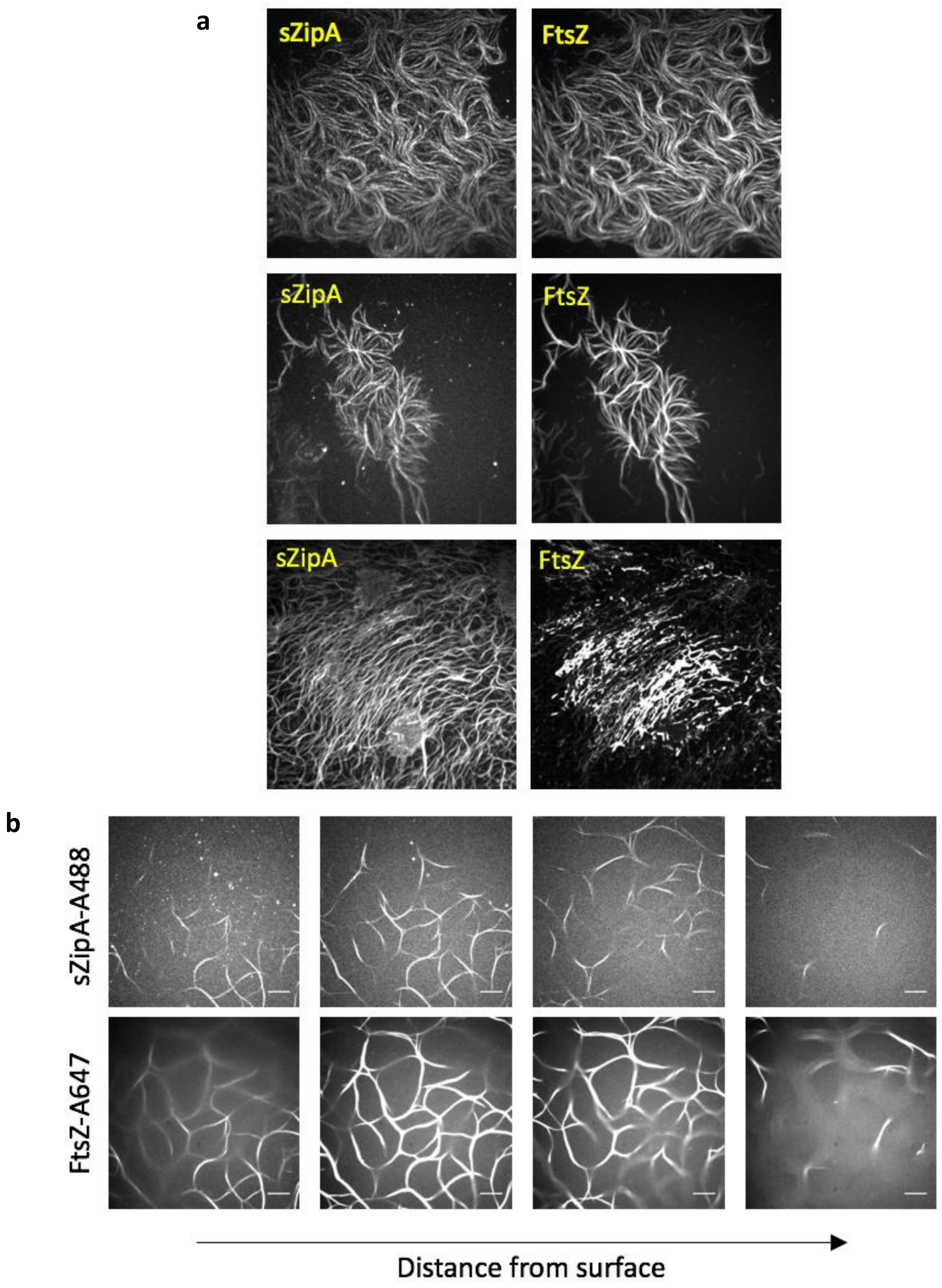


**Supplementary Fig. 7: Different phenotypes of sZipA-FtsZ co-filament networks in the presence of cell-free synthesized ZapA. a**, A solution containing purified FtsZ-A647, cell-free expressed ZapA from the *zapA*$_{opt}$ construct and additional 2 mM GTP, was incubated on top of an sZipA-A488-bound SLB. Images are from the FtsZ-A647 and sZipA-A488 channels, as annotated. **b**, Same conditions as in **a**. sZipA-FtsZ-ZapA co-filaments were abundantly found to point upwards the SLB and extend in the bulk phase. Both FtsZ-A647 and sZipA-A488 channels are displayed, as annotated. Scale bars are 10 µm.

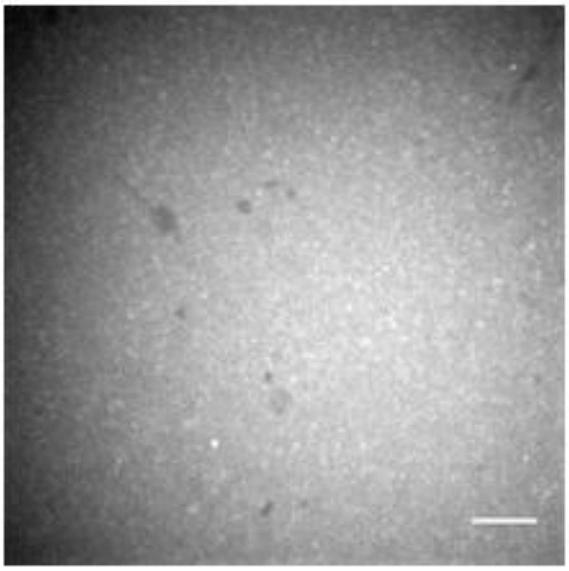

**Supplementary Fig. 8: Cell-free synthesized ZapA is unable to produce membrane-bound FtsZ filaments in the absence of ZipA and FtsA.** Concentration of purified FtsZ-A647 was 3 µM. Scale bar represents 10 µm.

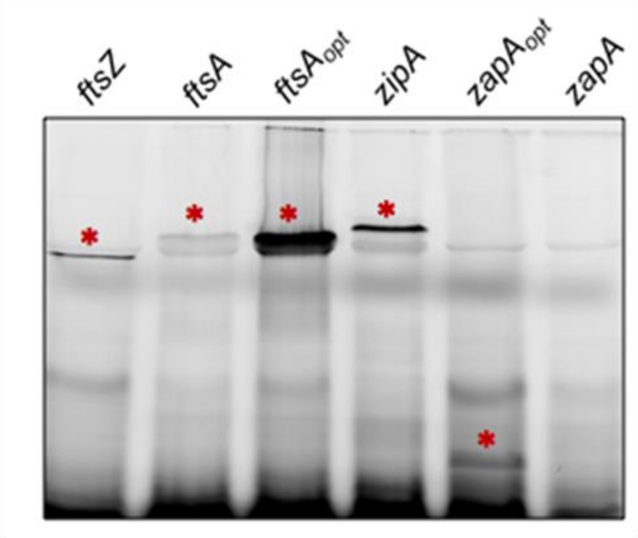


**Supplementary Fig. 9: Visualization of cell-free expressed proteins by SDS-PAGE imaging.** Single genes were expressed in PURE system reactions in the presence of the GreenLys reagent for co-translational labelling with a BODIPY-conjugated lysine residue. Translation products were analysed by fluorescence imaging of a 18% polyacrylamide gel. The bands depicted with an upper red star correspond to the full-length protein with expected molecular weight. The sequence-optimized constructs encoding FtsA and ZapA are named $ftsA_{opt}$ and $zapA_{opt}$, respectively. No specific band was clearly visible when the native sequence of the *zapA* gene was expressed, denoting a low protein yield.

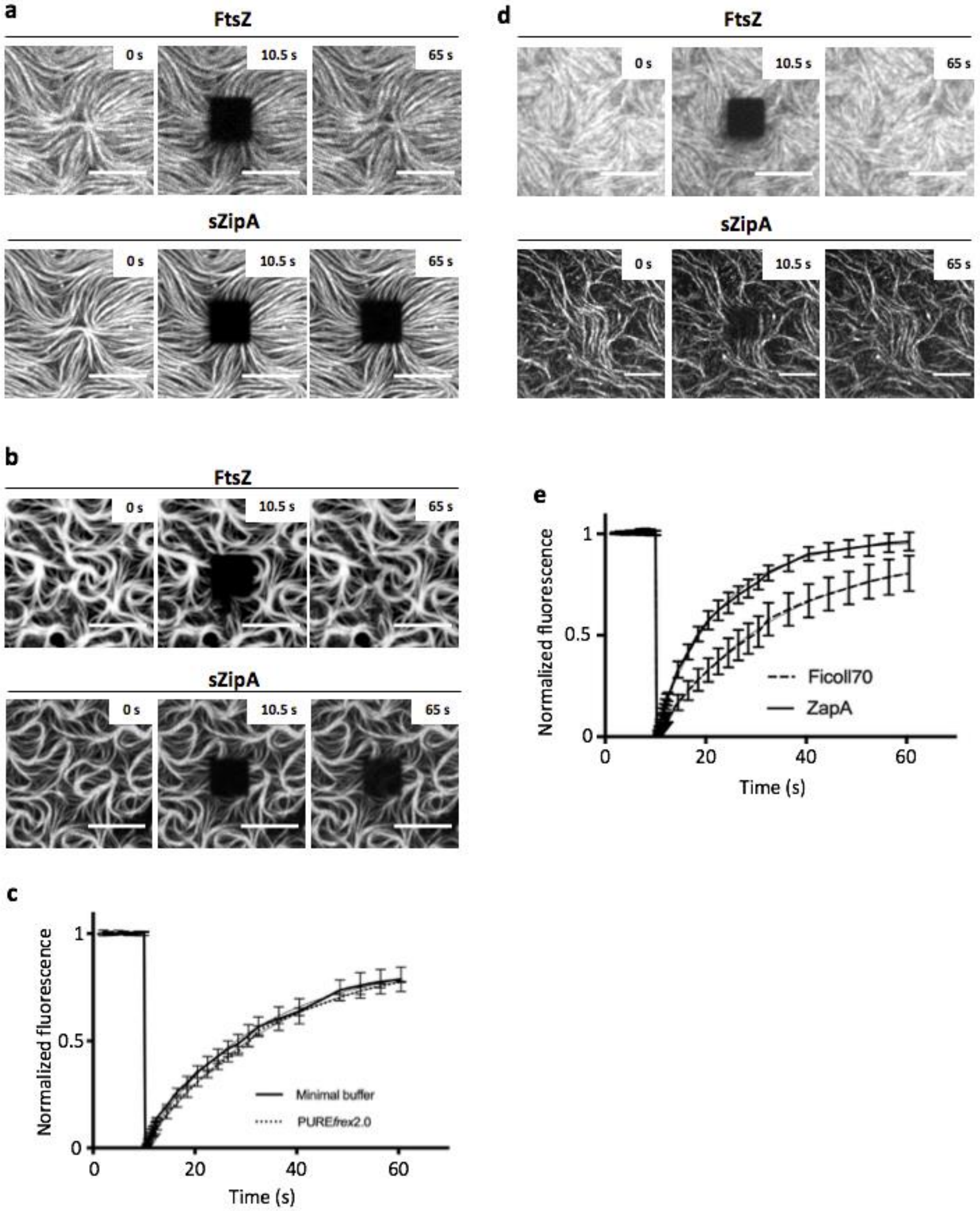


**Supplementary Fig. 10: Ficoll70 and cell-free expressed ZapA differently influence the dynamics of FtsZ and sZipA on SLBs. a**,**b** Time-lapsed fluorescence images of purified sZipA-A488 and FtsZ-A647 during FRAP experiments in minimal buffer (**a**) and PURE*frex*2.0 (**b**) containing Ficoll70. The conditions in **a** are identical as in the main text **Fig. 2c,d**. At 10.5 s a square area was bleached and fluorescence recovery was monitored in both the FtsZ and sZipA channels, as annotated. Scale bars indicate 10 µm. **c**, FRAP curves of FtsZ-A647 fluorescence according to the experiments shown in **a** and **b**. Data are average values from three FRAP experiments and error bars represent standard deviation values. Exponential fits are appended. No recovery of sZipA signal was observed in either buffer conditions indicating poor lateral diffusion and turnover of the membrane-bound sZipA. We found that the recovery time (or recovery half-time) of FtsZ is about 15.2 ± 1.4 s in minimal buffer and about 17.6 ± 1.3 s in PURE system background. This result indicates that the FtsZ turnover rate between the bulk and filaments is similar in the two media. **d**, Time-lapsed fluorescence images of purified sZipA-A488 and FtsZ-A647 during FRAP experiments with the cell-free expressed $zapA_{\mathrm{opt}}$ gene. The conditions are identical as in the main text **Fig. 2c**. At 10.5 s a square area was bleached and fluorescence recovery was monitored in both the FtsZ and sZipA channels, as annotated. Different fields of view are shown. Scale bars indicate 10 µm. **e**, FRAP curves of FtsZ-A647 fluorescence corresponding to the experiment shown in **d**. Data are average values from four FRAP experiments and error bars represent standard deviation values. Exponential fits are appended. FRAP experiments show that FtsZ-A647 monomers undergo twice faster exchange rate between filaments and bulk with ZapA expressed from the optimized or non-optimized construct than with Ficoll70 (~7.5 s vs. ~17 s).

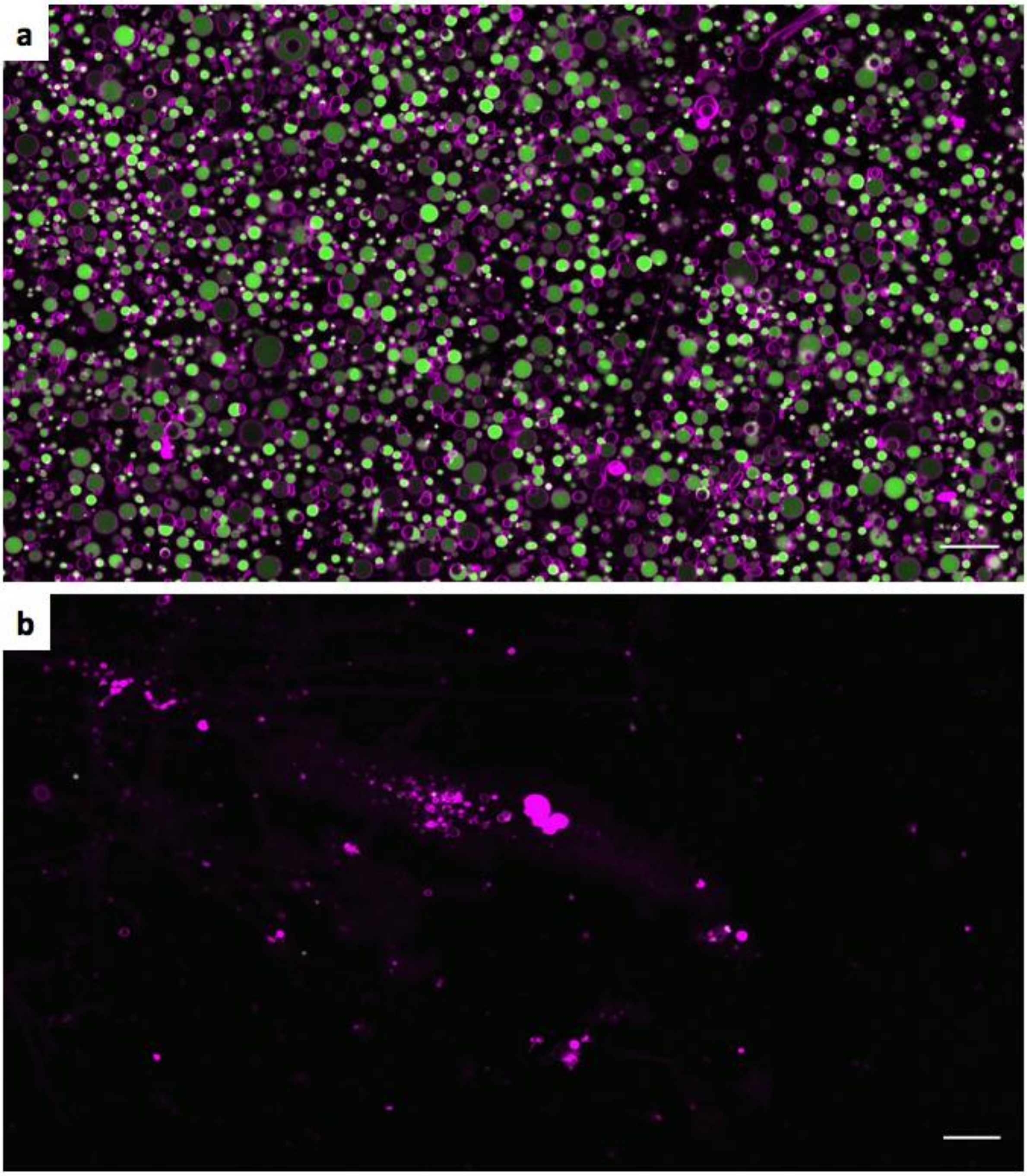


**Supplementary Fig. 11: Ficoll70 impairs formation of gene-expressing liposomes. a**, Fluorescence confocal image of liposomes expressing the *yfp* gene reporter. The same protocol was used to express cytoskeletal proteins, as shown in the main text **Fig. 3** and **4**. The membrane dye signal is coloured in magenta and the YFP fluorescence is colored in green. **b**, The presence of Ficoll70 in the swelling medium alters the production of liposomes. Scale bars represent 20 µm.

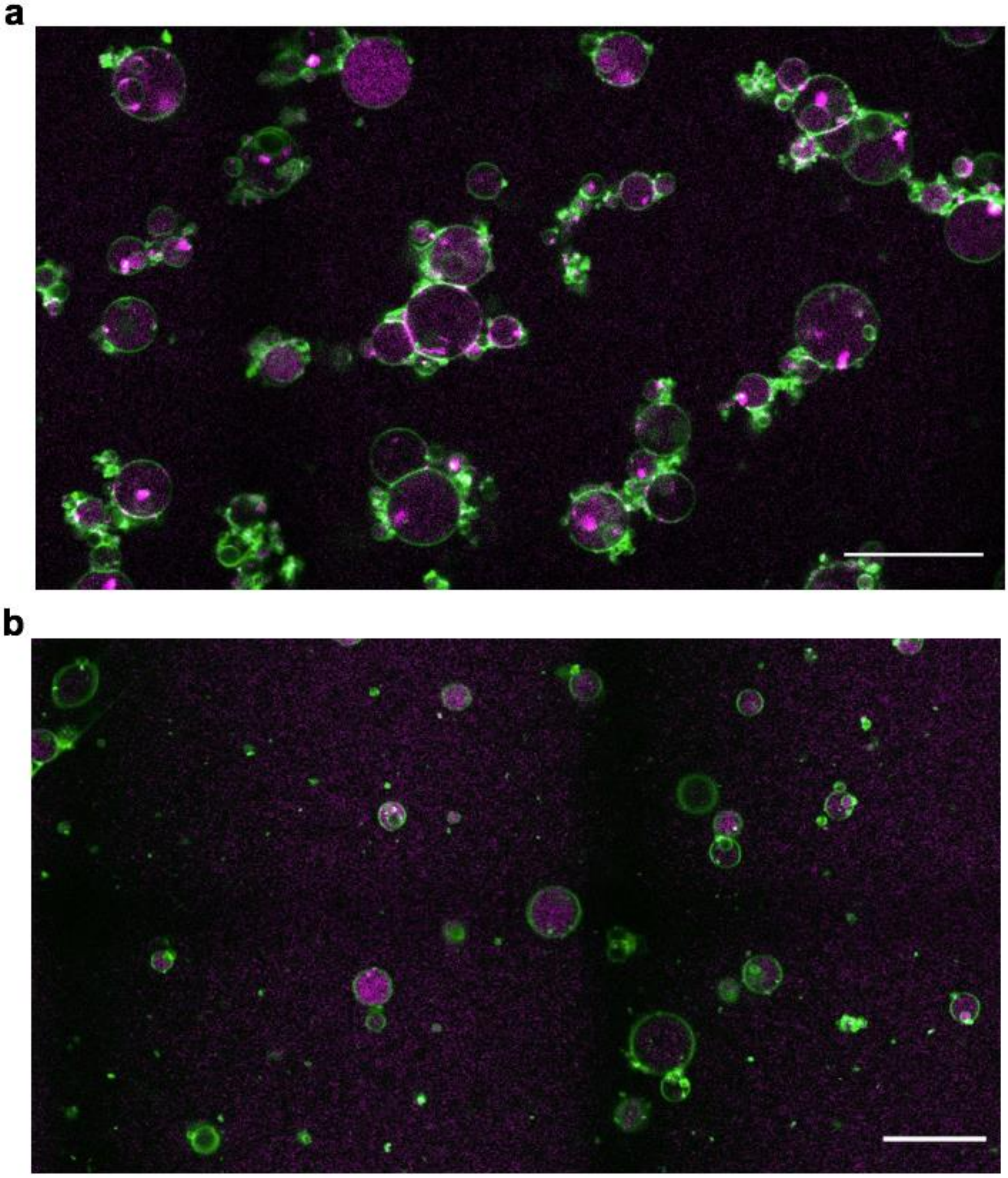


**Supplementary Fig. 12: Confocal fluorescence images of FtsZ-A647-containing liposomes with and without expressed FtsA.** **a**, Population of liposomes containing FtsA and purified FtsZ-A647 after 6 h expression of the *ftsA*$_{opt}$ construct. Agglutinated small liposomes can be seen around larger vesicles. **b**, Population of liposomes containing purified FtsZ-A647 but no *ftsA$_{opt}$* gene. The image was taken after 6 h incubation to reproduce the conditions for gene expression (**a**). The occurrence of small vesicles surrounding a big liposome is much lower than in the presence of in situ synthesized FtsA. Moreover, FtsZ-A647 is localized evenly in the lumen of vesicles. Membrane is coloured in green and FtsZ-A647 in red. Different imaging settings have been used compared to the image shown in **a**, explaining the difference in background signal in the FtsZ-A647channel. Scale bars represent 20 µm.

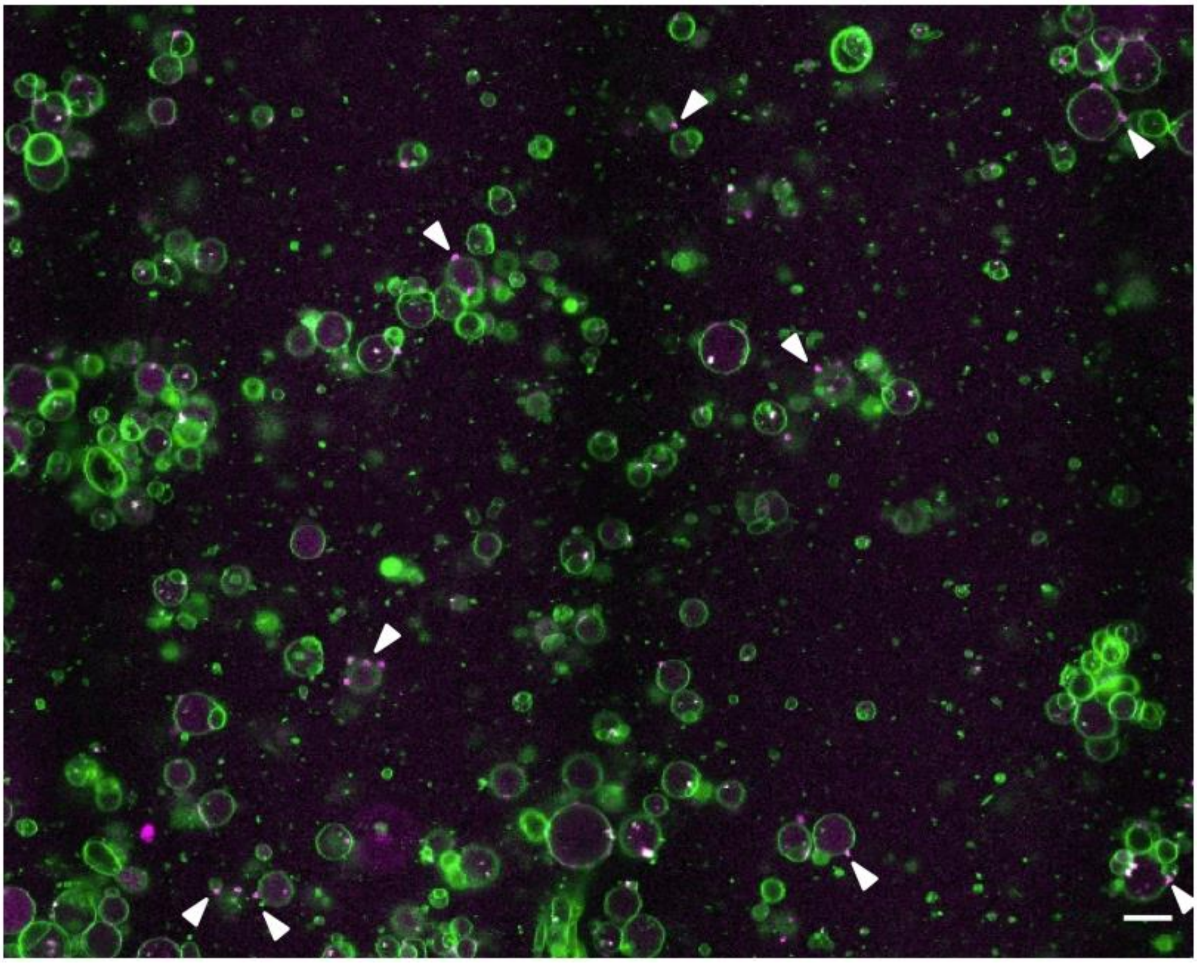

**Supplementary Fig. 13: Confocal fluorescence image of liposomes containing purified FtsZ-A647 and in situ expressed FtsA.** Experimental conditions are identical as in the main text **Fig. 3**. The image was taken after ~3.5 h incubation for gene expression. Fluorescence from the membrane dye is coloured in green and FtsZ-A647 signal is in magenta. Arrowheads point to liposomes exhibiting budding spots or constriction sites. Scale bar represents 20 µm.

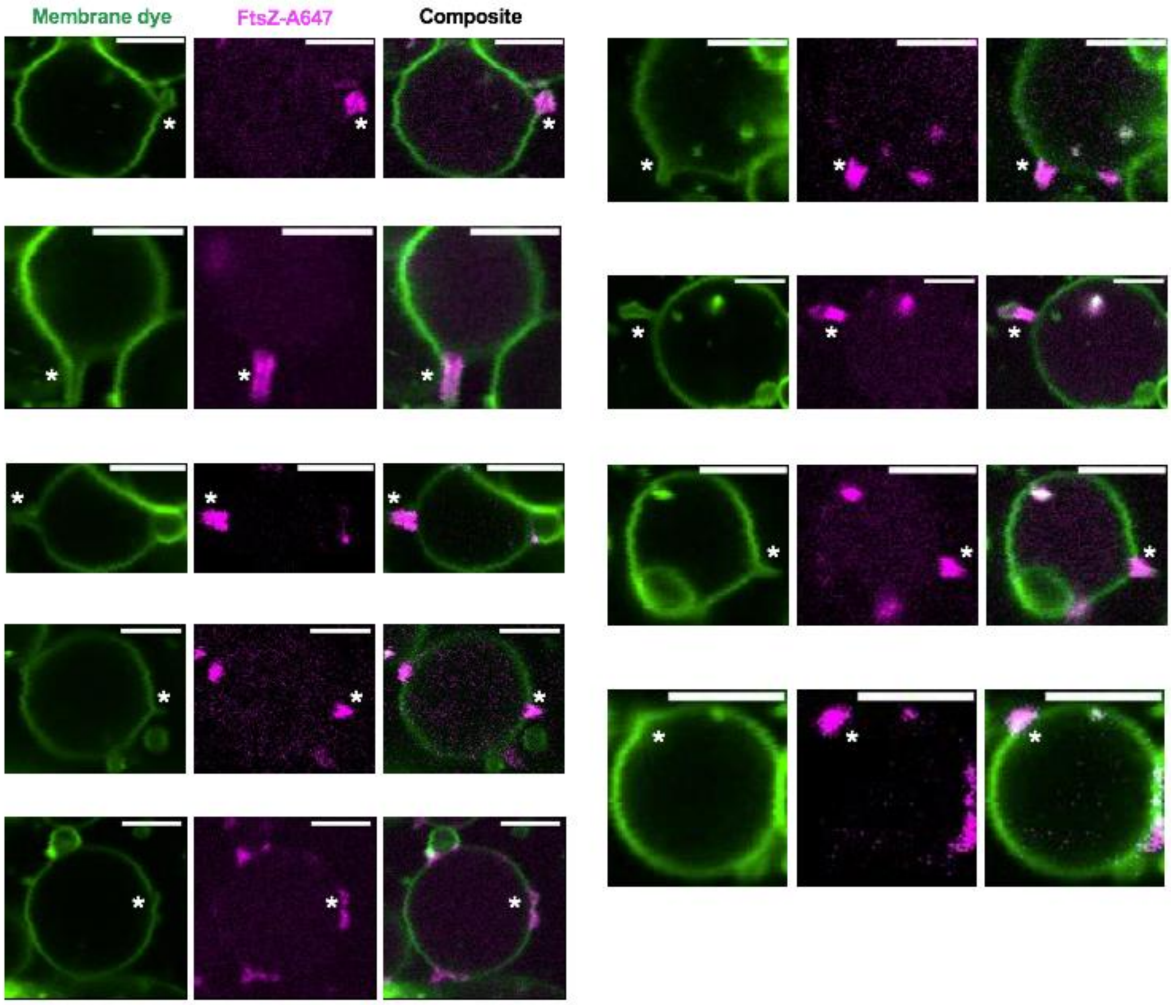


**Supplementary Fig. 14: Confocal fluorescence images of individual liposomes containing purified FtsZ-A647 and in situ expressed FtsA.** Experimental conditions are identical as in the main text **Fig. 3**. Fluorescence from the membrane dye is coloured in green and FtsZ-A647 signal is in magenta. The composite image is the overlay of the two channels. Asterisks indicate budding spots or constriction sites. Scale bars represent 5 µm.

### SUPPLEMENTARY REFERENCES


1. Martos, A. *et al.* Characterization of self-association and heteroassociation of bacterial cell division proteins FtsZ and ZipA in solution by composition gradient−static light scattering. *Biochemistry* **49**, 10780–10787 (2010).
2. Loose, M. & Mitchison, T. J. The bacterial cell division proteins FtsA and FtsZ self-organize into dynamic cytoskeletal patterns. *Nat. Cell Biol.* **16**, 38–46 (2014).
3. Wang, X. & Lutkenhaus, J. The FtsZ protein of *Bacillus subtilis* is localized at the division site and has GTPase activity that is dependent upon FtsZ concentration. *Mol. Microbiol.* **9**, 435–442 (1993).
4. Sossong, T. M., Brigham-Burke, M. R., Hensley, P. & Pearce, K. H. Self-activation of guanosine triphosphatase activity by oligomerization of the bacterial cell division protein FtsZ †. *Biochemistry* **38**, 14843–14850 (1999).
5. González, J. M. *et al.* Essential cell division protein FtsZ assembles into one monomer-thick ribbons under conditions resembling the crowded intracellular environment. *J. Biol. Chem.* **278**, 37664–71 (2003).
6. Martos, A. *et al.* FtsZ polymers tethered to the membrane by ZipA are susceptible to spatial regulation by Min waves. *Biophys. J.* **108**, 2371–2383 (2015).
7. Cabré, E. J. *et al.* Bacterial division proteins FtsZ and ZipA induce vesicle shrinkage and cell membrane invagination. *J. Biol. Chem.* **288**, 26625–34 (2013).
8. Furusato, T. *et al. De novo* synthesis of basal bacterial cell division proteins FtsZ, FtsA, and ZipA inside giant vesicles. *ACS Synth. Biol.* **7**, 953–961 (2018).
9. Rivas, G., Alfonso, C., Jiménez, M., Monterroso, B. & Zorrilla, S. Macromolecular interactions of the bacterial division FtsZ protein: from quantitative biochemistry and crowding to reconstructing minimal divisomes in the test tube. *Biophys. Rev.* **5**, 63–77 (2013).
